# Multitargeted Reduction of Inflammation and Atherosclerosis in *Tet2*-deficient CHIP via XPO1 Inhibition and Atf3 restoration

**DOI:** 10.1101/2025.06.12.658927

**Authors:** Nicole Prutsch, Amélie Vromman, Brittaney Leeper, Mengyu Chen, Julia Keating, Shuning He, Alla Berezovskaya, Siyang Ren, Mariana Janini Gomes, Eduardo J. Folco, Philipp J. Rauch, Prafulla C. Gokhale, Mark W. Zimmerman, Donna S. Neuberg, Brian J. Abraham, Peter Libby, A. Thomas Look

## Abstract

*TET2* is the second most frequently mutated gene in clonal hematopoiesis of indeterminate potential (CHIP), driving hematopoietic stem cell clonal expansion and increasing the risk of myeloid malignancies. Affected individuals often develop atherosclerotic cardiovascular disease, exacerbated by hyperinflammatory *TET2*-mutant macrophages. Here, we show that the XPO1 nuclear export inhibitor eltanexor significantly reduces atherosclerotic plaque formation in a mouse model of *Tet2*-mutant CHIP. In addition, we investigated the mechanisms and gene expression pathways that underlie the proinflammatory phenotype that characterizes *Tet2*-mutant CHIP. Single-cell CITE-seq identified increased expression of multiple proinflammatory mediators in *Tet2*-mutant macrophages and in non-hematopoietic cells of the aortic wall, which was reduced by eltanexor treatment. *Atf3*, which encodes a core transcriptional modulator of inflammation, occupies and regulates the largest enhancer in wild-type macrophages. *Tet2* loss diminished ATF3 binding to the regulatory loci of inflammatory mediators, which was restored upon XPO1 inhibition. These results provide new insights into drivers of heightened inflammation in *TET2*-mutant CHIP and highlight a novel therapeutic strategy for intervention.

## INTRODUCTION

Loss-of-function mutations of the epigenetic modifier TET2 in hematopoietic stem and progenitor cells (HPSC) frequently drive clonal hematopoiesis of indeterminate potential (CHIP) ^1–3^. These mutations result in the hypermethylation of key transcriptional enhancers that disrupt essential cellular processes such as self-renewal, proliferation, survival, and differentiation leading to clonal advantage of HSPCs in the bone marrow, which then give rise to *TET2*-mutant monocytes and macrophages in the periphery ^4,5^. Consequently, *TET2*-mutant CHIP links to an elevated risk of both hematological cancer and atherosclerotic and other cardiovascular and chronic conditions ^2,3,6–9^. The increased risk of severe atherosclerosis-related consequences in CHIP patients arises from a hyperinflammatory state exhibited by the mutant cells, particularly monocytes and macrophages, that display heightened production of pro-inflammatory chemokines and cytokines, including interleukins (IL)-1β and IL-6 ^2,10,11^, as well as the NLR3P inflammasome^12^. Several studies have shown that the escalated inflammation driven by the mature mutant myeloid cell progeny acts to reinforce the clonal advantage of mutant HSPCs in the bone marrow ^13–16^. However, the mechanisms that underlie the promotion of inflammation by TET2 loss that foster this re-enforcing cycle remain still poorly understood.

The consequences of *TET2*-mutant CHIP underscore the pressing need to identify drugs that specifically target *TET2*-mutant HSPCs and the differentiated cells they produce to reduce the number of mutant macrophages that have increased production of inflammatory mediators. The administration of IL-1β and IL6R blocking antibodies, as well as NLRP3 inflammasome inhibitors, to CHIP patients, may offer strategies to decrease the risk of cardiovascular morbidity in patients with *TET2*-mutant CHIP^11,17,18^. Agents like colchicine that suppress inflammation are also under consideration ^19^. However, an ideal therapeutic approach in CHIP would suppress both the growth of mutant HSPC clones in the bone marrow and the secretion of proinflammatory cytokines by *Tet2*-mutant macrophages, to limit both atherosclerotic cardiovascular disease and the progression to myeloid malignancy.

Our previous investigations identified efficacy of the nuclear export inhibitor eltanexor in reducing the number of *Tet2*-mutant HSPCs in zebrafish embryos and in murine *in vitro* colony formation assays ^20^. Here, we tested eltanexor in a mouse model of *Tet2*-mutant CHIP to test the hypothesis that eltanexor can simultaneously suppress *Tet2*-mutant clonal expansion in the bone marrow and mitigate atherogenesis. We used a combination of single-cell CITE-seq studies of cells extracted from the bone marrow and arterial wall, as well as enhancer profiling in murine bone marrow-derived macrophages, to identify the underlying mechanisms and gene expression pathways through which *Tet2* loss of function promotes clonal advantage, inflammation, and atherosclerotic plaque formation.

## RESULTS

### XPO1 inhibition significantly reduces atherosclerotic plaque formation in a murine model of *Tet2*-deficient CHIP

Previous studies have established that atherosclerosis-prone *Ldlr-/-* mice receiving *Tet2*-mutant bone marrow cells show increased atherosclerotic plaque formation when consuming a high-cholesterol diet, as compared to mice injected with wild-type bone marrow cells ^2,7,10^. Building on our published findings in which we identified selective activity of the nuclear export receptor exportin-1 (XPO1, CRM1) inhibitor eltanexor in killing *Tet2*-mutant HSPCs ^21^, we tested the activity of eltanexor in reducing *Tet2*-CHIP related atherosclerotic plaque formation (**Fig. 1**). To mimic the coexistence of both *Tet2+/+* wild-type and *Tet2+/-* heterozygous mutant hematopoietic cells that coexist in patients with CHIP, we transplanted 1 x 10^6^ *Tet2*+/-;CD45.1+ bone marrow donor cells along with equal numbers of *Tet2+/+*, CD45.2+ competitor bone marrow cells into lethally irradiated *Tet2*+/+;CD45.2+ *Ldlr-/-* recipients ^2,5^ (Group1, **Fig. 1a**). As a control, we transplanted *Tet2*+/+;CD45.2+ *Ldlr-/-* recipient mice with *Tet2*+/+;CD45.1+ bone marrow donor cells along with equal numbers of *Tet2+/+*;CD45.2+ cells (Group 2, **Fig. 1a**).

**Figure 1.**
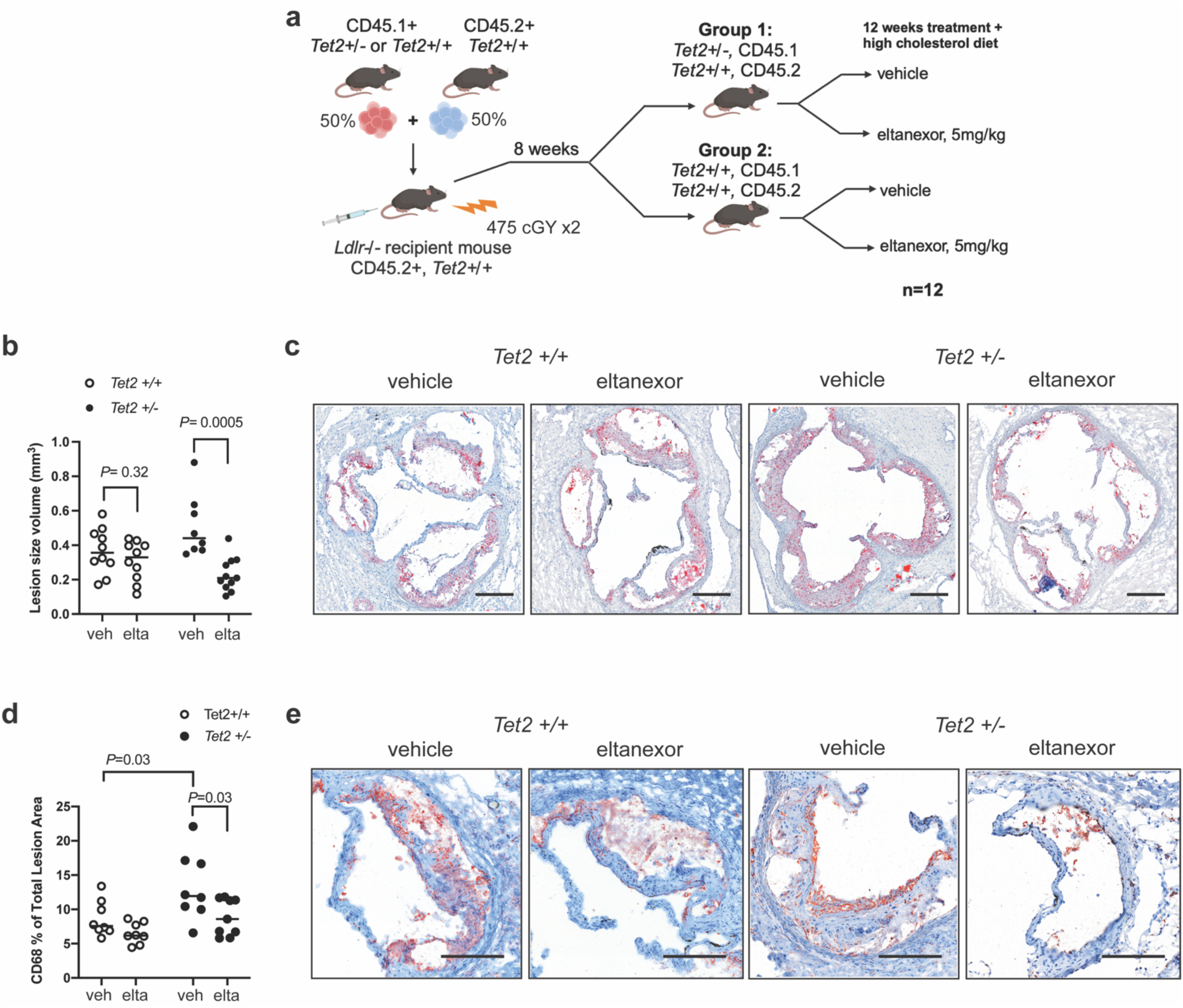
XPO1 inhibition significantly reduces atherosclerotic plaque formation in a murine model of *Tet2*-deficient CHIP. **a**, Female *Ldlr*-/-;*Tet2+/+*;CD45.2+ mice were transplanted with *Tet2+/-*;CD45.1+ (Group 1, n=24) or *Tet2+/+*;CD45.1+ (Group 2, n=24) 1x10e6 bone marrow cells in a 1:1 mix with 1x10e6 *Tet2+/+*;CD45.2+ normal *Tet2*+/+ bone marrow cells. After 8 weeks of bone marrow reconstitution, 12 mice from each Group were treated for 12 weeks with 5 mg/kg of eltanexor, and 12 mice with vehicle control and fed a high-cholesterol diet for the entire 12 weeks. **b**, Quantification of aortic root lesions after Oil red O staining in transplanted female Group 1 and Group 2 *Ldlr-/-* mice after 12 weeks of treatment and the high-cholesterol diet. P-values obtained by the Wilcoxon rank-sum test. **c**, Representative images of aortic root sections stained with Oil red O and shown at 20x magnification (scale bar = 200mm). **d**, Quantification of CD68+ macrophage infiltration in the aortic root lesions from the 4 subgroups of n=12 transplanted mice treated as indicated in Panel **a**. P-values obtained by the Wilcoxon rank-sum test. **e**, Representative images of aortic root sections stained by IHC with anti-CD68 antibody, shown at 33x magnification (scale bar = 200mm).

Eight weeks post-transplantation, mice with successful engraftment were randomized into treatment arms of 12 mice, each receiving either eltanexor (5 mg/kg QD, 5 days/week) or vehicle for 12 weeks by oral gavage. In parallel, all mice received a high-cholesterol diet to promote atherosclerosis. After 12 weeks of treatment and high-cholesterol diet, we euthanized the mice to quantify the extent of atherosclerotic plaque formation within the aortic roots. Oil red O (ORO) staining measured the volume of atherosclerotic plaque over serial sections of the aortic roots of the *Ldlr-/-* mice (**Fig.1** and Methods). Compared to vehicle-treated mice receiving control *Tet2*+/+;CD45.1+ bone marrow cells (Group 2), vehicle-treated recipients of *Tet2*+/-;CD45.1+ bone marrow cells (Group 1) showed a calculated 3D lesion size in the aortic root that was a median 23.6% percent larger (0.356 vs. 0.440 mm^3^) after 12 weeks on the high-cholesterol diet (**Fig. 1b,c**). Consistent with our previous results showing selective activity of eltanexor against *Tet2*-mutant HSPCs *in vivo* in zebrafish embryos and in *in vitro* murine colony forming assays ^21^, 12 weeks of treatment with eltanexor significantly reduced atherosclerotic plaque formation compared to vehicle control in the aortic roots of *Ldlr-/-* mice receiving *Tet2+/-* bone marrow cells (Group 1), with a median lesion volume that was 52.3% smaller (0.440 vs. 0.210 mm^3^, *P*=0.0005) than in vehicle-treated mice (**Fig. 1b,c**). By contrast, eltanexor did not reduce atherosclerotic plaque formation in the aortic roots of *Ldlr-/-* mice injected with *Tet2^+/+^* control bone marrow (Group 2, 0.356 vs. 0.33 mm^3^, *P*=0.32), indicating selective activity of eltanexor in the mice receiving the *Tet2*-mutant bone marrow cells (**Fig. 1b,c**).

Previous studies demonstrated that *Tet2* loss-of-function promotes monocyte accumulation in the arterial wall and alters macrophage function in plaques to enhance atherosclerosis^1,2,7,15,22^. We therefore performed immunohistochemistry with the macrophage marker CD68 on aortic root sections to analyze the percentage of macrophages in the atherosclerotic plaques in each group, both with and without eltanexor treatment. Compared to *Ldlr-/-* mice receiving control *Tet2*+/+;CD45.1+ bone marrow cells (Group 2), recipients of *Tet2*+/-;CD45.1+ bone marrow cells (Group 1) showed a median percentage of macrophages in the atherosclerotic lesions that was 1.6-fold larger (7.6% versus 11.9%) after 12 weeks on the high-cholesterol diet (*P*=0.03) **Fig. 1d,e**). Eltanexor treatment in Group 1 *Ldlr-/-* mice injected with *Tet2+/-* bone marrow cells significantly reduced the CD68-expressing macrophages in the atherosclerotic plaques, with a median percentage that was 1.4-fold smaller (11.9% versus 8.6%, *P*=0.03) than in vehicle-treated Group 1 *Ldlr-/-* mice (**Fig. 1d,e**). Eltanexor treatment did not reduce the percentage of CD68+ macrophages in the atherosclerotic lesions of Group 2 control mice injected with *Tet2+/+* CD45.1+ control bone marrow cells (**Fig. 1d,e**). The selective activity of eltanexor in *Tet2*-mutant cells was also evident in the spleen, which showed a significant reduction due to eltanexor treatment in the spleen weight of Group1 *Ldlr-/-* mice injected with *Tet2+/-* bone marrow (*P*=0.0002), but not in Group2 *Ldlr-/-* mice injected with *Tet2+/+* control bone marrow cells (**Extended Data Fig. 1**). Importantly, while none of the eltanexor-treated mice lost weight during the course of this experiment, *Ldlr-/-* mice in both Group1 and Group 2 treated with eltanexor did not gain as much weight as mice in the vehicle-treated control groups (**Extended Data Fig. 2a**). Moreover, plasma cholesterol levels of eltanexor-treated mice were significantly lower than in vehicle-treated mice (**Extended Data Fig. 2b**). However, significant reduction in atherosclerotic plaques, macrophage content, and spleen weight was observed exclusively in mice injected with *Tet2+/-* bone marrow cells, but not in those injected with Tet2+/+ control marrow cells (**Fig. 1b-e, Extended Data Fig. 2**). Thus, our results demonstrate selective activity of eltanexor in targeting *Tet2+/-* mutant hematopoietic cells to decrease atherosclerotic plaque formation and macrophage infiltration in mice injected with *Tet2+/-* mutant bone marrow cells (**Fig. 1b-e**).

### XPO1 inhibition reduces *Tet2+/-* mutant HSCs and CMP numbers and alters the expression of genes associated with self-renewal and inflammation in the bone marrow

*Tet2* loss in CHIP forms a self-reinforcing circuit of increased pro-inflammatory cytokine production in *Tet2*-mutant macrophages that, in turn, stimulates the growth of *Tet2*-mutant HSPCs in the bone marrow to promote their clonal expansion ^13,15,16^. To determine whether the reduction of atherosclerotic plaques is associated with an effect of eltanexor on the HSPCs of the *Tet2*-mutant clone in the bone marrow, we analyzed RNA expression patterns by single-cell CITE-seq in Lin-, Ska+, Kit+ (LSK)-sorted murine bone marrow cells of *Ldlr-/-* mice transplanted with *Tet2* +/- or *Tet2* +/+ bone marrow and treated for 12 weeks with eltanexor or vehicle (**Fig. 2a top**). Within these LSK cells, we identified 16 different UMAP clusters of stem and progenitor cells based on the most differentially expressed genes (**Fig. 2b, Supplementary Table 1**). We additionally used antibody detection of cell surface protein expression to distinguish between the LSK cells expressing CD45.1 or CD45.2 within the Group 1 and Group 2 competitive repopulation experiments (**Fig. 1a**, **Fig. 2a bottom**). Group 1 mice showed approximately twice as many *Tet2+/-*;CD45.1+ cells compared to *Tet2+/+;*CD45.2+ cells in each of the 16 vehicle-treated cell clusters (67% versus 33% respectively, see frequency chart in **Fig. 2b top**), reflecting the clonal advantage of the *Tet2+/-*;CD45.1+ mutant LSK-sorted bone marrow cells. Interestingly, in Group 2, we observed an increase in the Tet2+/+; CD45.2+ cell fraction as compared to the *Tet2+/+*;CD45.1+ cell fraction within the vehicle-treated mice (82.2% versus 17.8% respectively, see frequency chart in **Extended Data Fig. 3**), indicating that the *Tet2+/+*;CD45.2+ cells from the residual bone marrow cells of the irradiated CD45.2+ host mice contribute to the *Tet2+/+*;CD45.2+ population in these experiments. The excess of CD45.2+ cells in Group 2 control mice suggests that the clonal dominance of the *Tet2+/-*;CD45.1+ mutant LSK-sorted bone marrow cells has an even greater impact than the observed difference in vehicle-treated Group 1 mice (**Fig. 2b top**). While eltanexor treatment did not reduce the clonal advantage of the overall *Tet2+/-*;CD45.1+ LSK fraction in Group 1 mice (**Fig. 2b top**), we observed a slight reduction of *Tet2+/-*;CD45.1+ HSCs in Cluster 1 as compared to Tet2+/+;CD45.2 HSCs (8.9% vs. 9.5% respectively) and a strong reduction of *Tet2+/-*;CD45.1+ common myeloid progenitors (CMP) in Cluster 3 as compared to *Tet2+/+;*CD45.2 CMPs (4.9% to 8.5% respectively) (**Fig. 2b top**, **Fig. 2c**). By contrast, eltanexor treatment increased the fraction of *Tet2+/-*;CD45.1+ multipotent progenitor cells (MPP) in Cluster 0, B-cell and dendritic cell progenitors in Cluster 4, monocyte-macrophage progenitors (MMP) in Cluster 8, and B- cell progenitors in Clusters 10 and 12 (**Fig. 2b top**).

**Figure 2.**
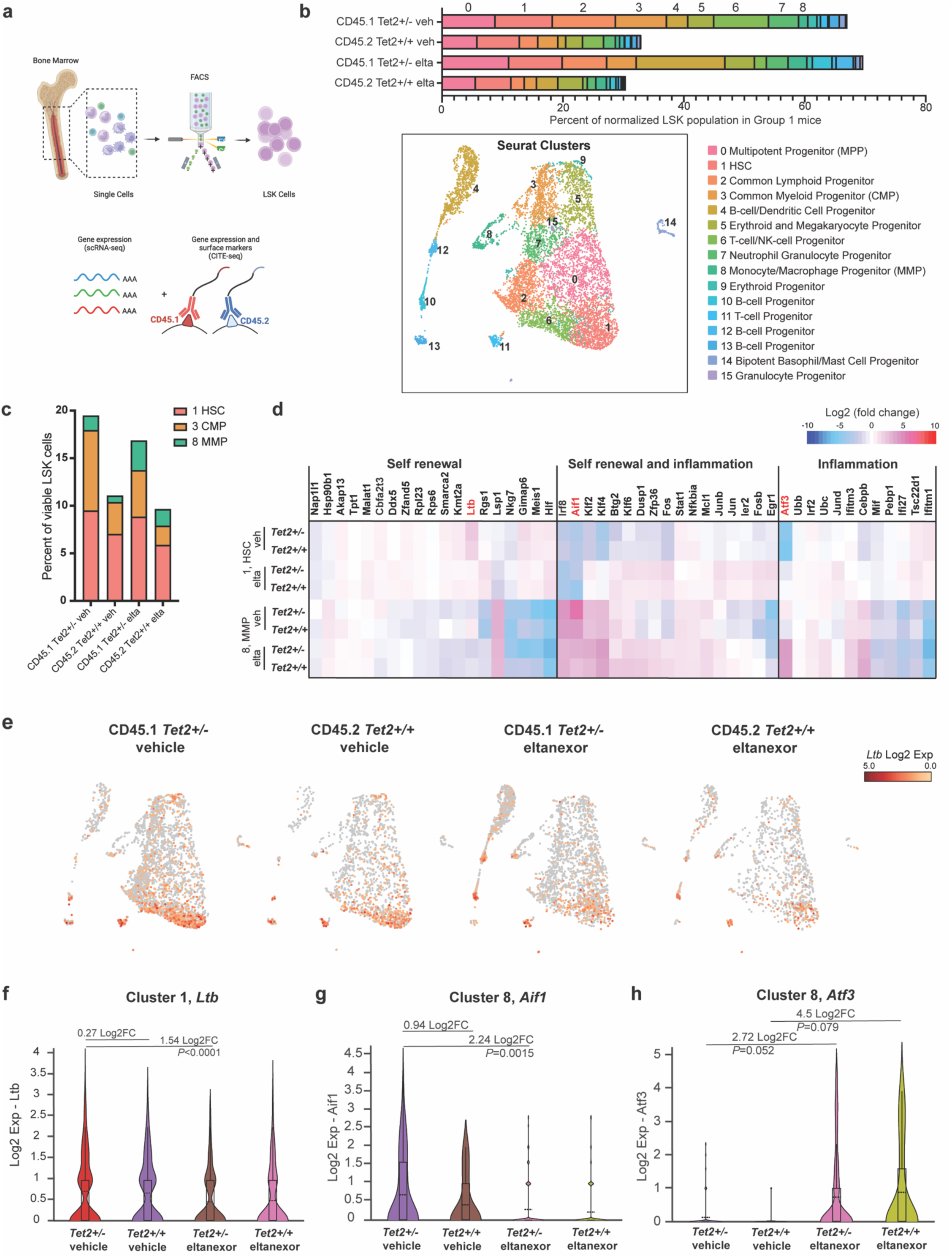
XPO1 inhibition reduces *Tet2+/-* mutant HSCs and CMP numbers and alters the expression of genes associated with self-renewal and inflammation in the bone marrow. **a,** Analysis of Group 1 mice (competitively transplanted with Tet2+/-; CD45.1+ as indicated in Figure 1a). After 12 weeks on a high cholesterol diet and treatment with either 5 mg/kg of eltanexor daily by oral gavage or vehicle control, BM cells from both femurs and spine of four mice treated with eltanexor and 4 mice treated with vehicle control were harvested and sorted for LSK cells by FACS. Viable cells were stained on the cell surface with CD45.1- and CD45.2-specific oligo-labeled antibodies and then analyzed by 10X single-cell RNA (scRNA) sequencing. **b**, UMAP of 9343 cells representing 16 clusters based on expression levels of the most differentially expressed genes (bottom panel) and a frequency bar chart representing their proportions in CD45.1+ and CD45.2+ cell populations of vehicle or eltanexor treated mice (top panel). Cells in each cluster were identified based on the most differentially expressed genes. Clusters include (0) multipotent progenitors (MPP), (1) hematopoietic stem cells (HSC), (2) common lymphoid progenitors, (3) common myeloid progenitors (CMP), (4) B-cell/dendritic cell progenitors, (5) erythroid and megakaryocyte progenitors, (6) T-cell/NK-cell progenitors, (7) neutrophil/granulocyte progenitors, (8) monocyte/macrophage progenitors (MMP), (9) erythroid progenitors, (10, 12, 13) B-cell progenitors, (11) T-cell progenitors, (14) bipotent basophil/mast cell progenitors, and (15) granulocyte progenitors. **c**, Frequency bar chart comparing the proportions of HSCs, CMPs and MMPs between the CD45.1+ Tet2+/- and CD45.2+ *Tet2+/+* cell populations in Group 1 mice treated with vehicle or eltanexor **d**, Heatmap of differentially expressed genes (DEGs) in *Tet2+/-*;CD45.1+ relative to *Tet2+/+*;CD45.2+ for HSCs and MMPs with and without eltanexor treatment. The genes shown are focusing on genes associated with “Self-renewal”, “Self-renewal and inflammation”, or “Inflammation” among the top 100 differentially expressed genes ^23^. **e,** UMAP depicting *Ltb* gene expression in HSCs from *Tet2+/-*;CD45.1+ and *Tet2+/+*;CD45.2+ cell populations in mice treated with vehicle or eltanexor. **f-h**, Violin plots of scRNA-seq data from *Tet2+/-*;CD45.1+ and *Tet2+/+*;CD45.2+ cell populations in mice treated with vehicle or eltanexor for (f) *Calr* in HSC (cluster 1), (g) *Aif1* and (h) *Atf3* in MMP (cluster 8).

Next, we analyzed the overall differences in gene expression due to *Tet2* mutation and eltanexor treatment across each of the 16 clusters. We particularly focused on Cluster 1 HSCs regarding mechanisms of altered self-renewal and Cluster 8 MMPs regarding altered inflammation. Expression of genes associated with self-renewal and inflammation^23^ in Cluster 1 of vehicle-treated mice exhibited a modest increase in Lymphotoxin beta (*Ltb*) expression in *Tet2+/-*;CD45.1+ compared to *Tet2+/+*;CD45.2 cells (**Fig. 2d,e,f**). Ltb is a type II membrane signaling protein of the TNF family that regulates self-renewal and differentiation of hematopoietic and leukemia stem cells and induces the inflammatory response system ^24^. Importantly, expression of *Ltb,* which is an XPO1 mRNA cargo ^25^, fell significantly with eltanexor treatment in *Tet2+/-*,CD45.1+ (P<0.0001) but not in *Tet2+/+*,CD45.2 HSCs (**Fig. 2d,e,f**). In the MMPs of Cluster 8, we found that allograft Inflammatory factor 1 (*Aif1*), a gene that is linked with the activation of macrophages and was initially identified in atherosclerotic plaques ^26–28^, rose strongly in *Tet2+/-*;CD45.1+ MMPs and fell significantly with eltanexor treatment in these cells (*P*=0.0015) but again, not in *Tet2+/+*,CD45.2 MMPs (**Fig. 2d, g**). Interestingly, activator of transcription factor 3 (*Atf3*), a member of the ATF/cAMP-response element binding protein (CREB) family of transcription factors, which inhibits the expression of IL1beta and several other inflammatory cytokines ^29,30^ rose in MMPs after eltanexor treatment (**Fig. 2d, h**). Taken together, these data indicate that eltanexor specifically reduces the clonal advantage of *Tet2*-mutant CMPs and alters the expression of key genes involved in self-renewal, proliferation, and inflammation of HSCs and MMPs.

### XPO1 inhibition reduces both *Tet2+/-* proinflammatory macrophage numbers within aortic arches and their expression levels of proinflammatory cytokines and chemokines

To determine the mechanisms by which eltanexor reduces formation of atherosclerotic plaque and the percentage of CD68+ *Tet2+/-* macrophages in lesions, we performed single-cell CITE-seq analysis on cells dissected from atherosclerotic plaque of the aortic arches of the *Ldlr-/-* mice (pool of 4 mice per treatment group) after 12 weeks of treatment (**Fig. 3a, top**). Based on the most differentially expressed genes, we identified 11 different cell type clusters using UMAP analysis (**Supplementary Table 2, Fig. 3b**). In the single-cell RNAseq experiment, we additionally used oligo-labeled cell surface antibodies to identify CD45.1+ or CD45.2+ hematopoietic cells, and non-hematopoietic CD45-negative stromal cells within the aortic arches of the transplanted mice (**Fig. 3a bottom, Fig. 3b**). The CD45-positive hematopoietic clusters included M1 (Cluster 2) and M2 macrophages (Cluster 5), monocytes (Cluster 8), cytotoxic T-cells (Cluster 4), double-positive T cells (Cluster 10), and B-cells (Cluster 7) (**Fig. 3b, right, Extended Data Fig. 4, 5**). The CD45-negative, non-hematopoietic cell types included myofibroblasts (Cluster 0), cardiomyocytes (Cluster 1), fibroblasts (Cluster 3), smooth muscle cells (Cluster 6), and endothelial cells (Cluster 9) (**Figure 3b, left**).

**Figure 3.**
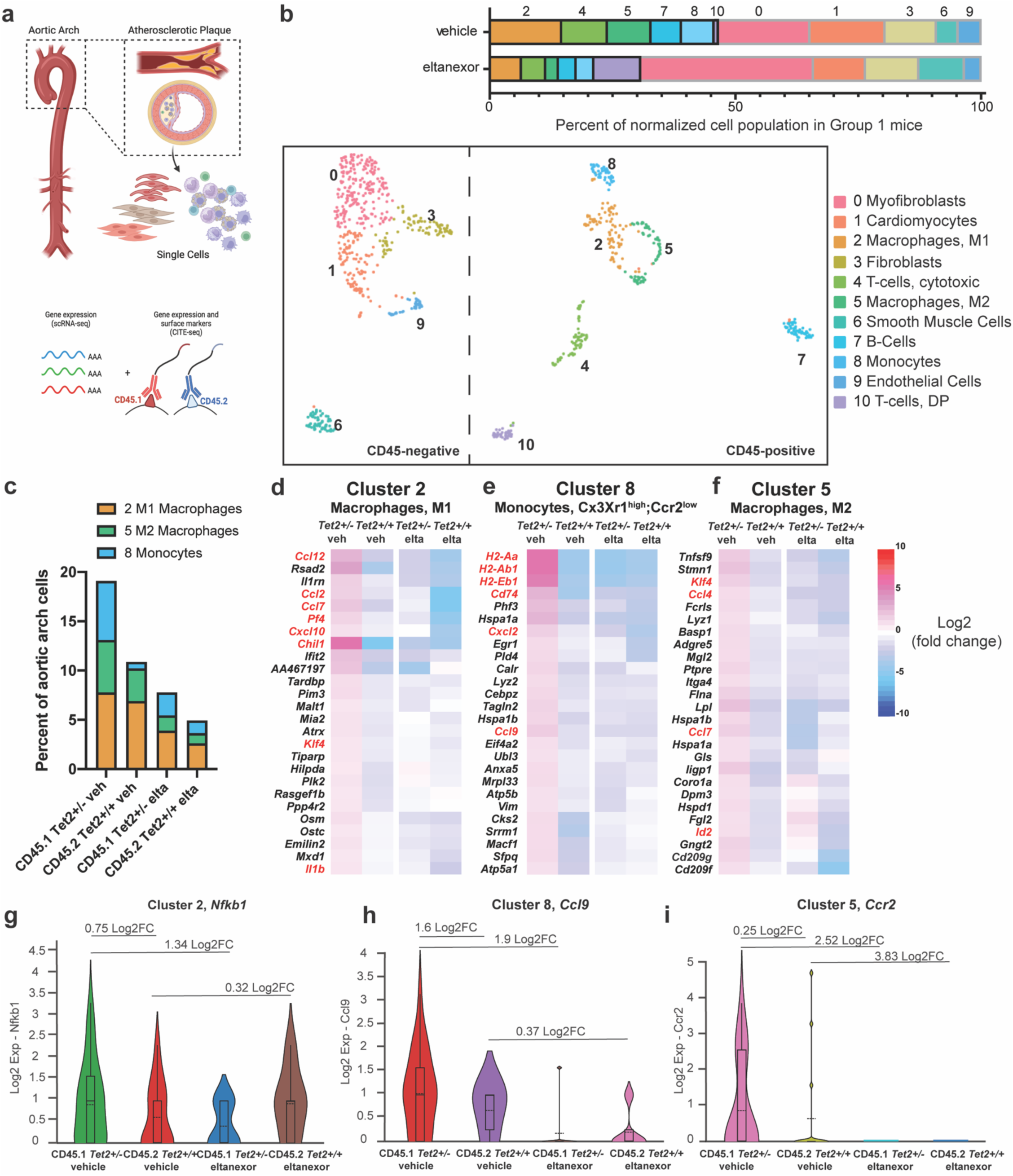
XPO1 inhibition by eltanexor reduces both *Tet2+/-* proinflammatory macrophage numbers within the aortic arches and their expression levels of proinflammatory cytokines and chemokines. **a,** Analysis of Group 1 mice (competitively transplanted with Tet2+/-; CD45.1+ as indicated in Figure 1a). After 12 weeks on a high cholesterol diet and treatment with either 5 mg/kg of eltanexor daily by oral gavage or vehicle control daily by oral gavage, aortic arches from four eltanexor-treated mice and four vehicle-treated mice were dissected and digested to obtain single cells. Cells were stained with CD45.1 and CD45.2 oligo-labeled, antibodies and analyzed by 10X single-cell RNA (scRNA) sequencing. **b**, UMAP of 1284 cells representing 11 clusters based on expression levels of the most differentially expressed genes and bar graph representing their proportions. These UMAP clusters included non-hematopoietic CD45-negative cell clusters contributed by the recipient mouse: (0) myofibroblasts, (1) cardiomyocytes, (3) fibroblasts, (6) smooth muscle cells, and (9) endothelial cells. Hematopoietic CD45-positive cell clusters were (2) M1 macrophages, (4) cytotoxic T-cells, (5) M2 macrophages, (7) B-cells, (8) monocytes, and (10) double-positive (DP) T-cells. **c**, Frequency bar chart comparing the proportions of M1 macrophages, M2 macrophages and monocytes divided into *Tet2+/-*;CD45.1+ and *Tet2+/+*;CD45.2+ cell populations in mice treated with vehicle or eltanexor. **d-f**, Heatmap of differentially expressed genes in *Tet2+/-*;CD45.1+ relative to *Tet2+/+*;CD45.2+ cells with and without eltanexor treatment for (d) M1 macrophages (cluster 2), (e) monocytes (Cluster 8) and (f) M2 macrophages (Cluster 5). **g-i**, Violin plots of scRNA-seq data from *Tet2+/-*;CD45.1+ and *Tet2+/+*;CD45.2+ cell populations in mice treated with vehicle or eltanexor for (g) *Nfkb1* in M1 macrophages (cluster 2), (h) *Ccl9* in monocytes (cluster 8) and (i) *Ccr2* in M2 macrophages (cluster 5).

Overall, in Group 1 mice reconstituted with *Tet2+/-*;CD45.1+ bone marrow cells, eltanexor treatment reduced the percentage of hematopoietic cells by half when compared to vehicle treated mice (65.2% vs. 30.7% respectively, see frequency chart in **Fig. 3b**, **top**). Specifically, among the six hematopoietic clusters, M1 macrophages in Cluster 2 fell from 14.6% to 6.4%, M2 macrophages in Cluster 5 from 8.9 to 6.2%, monocytes in Cluster 8 from 6.7% to 3.6%, cytotoxic T-cells in Cluster 4 from 9.3% to 4.9%, and B-cells in Cluster 7 from 6.2% to 3.6% (**Fig. 3b**, **top**). Cluster 10 contained double-positive (CD4+/CD8+) T-cells as part of the hematopoietic population. However, relatively few of these cells (less than 1%) localized in the arterial wall of mice treated with vehicle control (**Fig. 3b**, **top**). We and others have observed that peripheral double-positive T cells represent a small subpopulation in mice after bone marrow transplantation^31^. Thus, although these double-positive T-cells increased by nearly 10-fold in eltanexor-treated mice (0.9% to 9.6%), these cells are limited to transplanted mice, and have unclear functional relevance.

We next focused on *Tet2+/-*;CD45.1+ and *Tet2+/+*;CD45.2+ cell subpopulations within the macrophage and monocyte clusters of Group 1 mice (**Fig. 3c**). We observed the most dramatic differences in monocytes (Cluster 8, blue) of vehicle-treated mice, with a 9-fold increase in the percentage of *Tet2*+/-;CD45.1+ monocytes compared to *Tet2+/+*;CD45.2+ monocytes (6% vs. 0.7%), reflecting the increased clonal advantage and invasiveness of the *Tet2*+/-;CD45.1+ monocytes (**Fig. 3c**). Eltanexor treatment dramatically reduced the *Tet2*+/-;CD45.1+ Cluster 8 monocytes from 6% to 2.3%, bringing their percentage closer to that of *Tet2*+/+;CD45.2+ wild- type Cluster 8 monocytes (**Fig. 3c).** Cluster 2, which encompasses M1 macrophages, had nearly equal percentages of *Tet2*+/-;CD45.1+ mutant and *Tet2*+/+;CD45.2+ wild-type M1 macrophages (7.8% vs. 6.9%, respectively) in mice treated with vehicle control. Eltanexor reduced the percentages of M1 macrophages of both genotypes, reducing the *Tet2+/-*;CD45.1+ M1 macrophages to 3.9% and the *Tet2+/+*;CD45.2 macrophages to 2.6% (**Fig. 3c)**. Similarly, eltanexor markedly reduced the percentages of both *Tet2+/-*;CD45.1+ and *Tet2+/+*;CD45.2+ M2 macrophages in cluster 5 (from 5.3% to 1.5% and from 3.3% to 1%, respectively) in the arterial walls of the transplanted Group 1 mice (**Fig. 3c)**.

Next, we compared single-cell gene expression profiles between the three different subpopulations of monocytes (Cluster 8) and macrophages (Clusters 2 and 5) (**Fig. 3 d-f**). First, we compared the most differentially expressed genes of the *Tet2+/-*;CD45.1+ with the *Tet2+/+*;CD45.2+ wild-type Cluster 2 M1 proinflammatory macrophages. The vehicle-treated mice displayed a strong proinflammatory gene expression profile in *Tet2+/-*;CD45.1+ macrophages with increased expression of secreted chemokines and cytokines, including *Ccl2*, *Ccl7*, *Ccl12*, as well as *Pf4* and *Il1b* compared to *Tet2+/+*;CD45.2+ macrophages (**Fig. 3d**). Moreover, the expression levels of the chemokine receptor *Cxcl10*, the transcription factor Kruppel-like factor 4 (*Klf4*), a Yamanaka factor ^32^ and mediator of pro-inflammatory signaling in macrophages ^33^, and Chitinase 3-like 1 (*Chil1*), a secreted inflammatory mediator of macrophages ^34^, were also increased in the vehicle-treated *Tet2+/-*;CD45.1+ mutant macrophages. Importantly, eltanexor treatment resulted in decreased expression levels, not only of *Il1b,* but each of the upregulated cytokines and chemokines in *Tet2+/-*;CD45.1+ M1 macrophages to the levels found in *Tet2+/+*;CD45.2+ control M1 macrophages (**Fig. 3d** and **Extended Data Fig. 6a**). *Nfkb1,* a pivotal mediator of inflammatory responses that regulates numerous pro-inflammatory cytokines including TNFα, IL1b, and IL6^35–37^, also rose in Cluster 2 *Tet2+/-*;CD45.1+ mutant M1 macrophages and fell with eltanexor treatment, likely because it is an XPO1 cargo^25^ (**Fig. 3g**). These results indicate that eltanexor not only reduces the relative numbers of *Tet2+/-*;CD45.1+ M1 macrophages in the arterial wall, but also globally limits their production of proinflammatory mediators.

Next, we focused on Cluster 8, which we identified as *Cx3Cr1^high^, Ccr2^low^* monocytes (**Fig. 3e, Extended Data Fig. 4**). As in the Cluster 2 macrophages, *Tet2+/-*;CD45.1+ monocytes had an accentuated proinflammatory profile compared to *Tet2+/+*;CD45.2+ control monocytes as gauged by increased expression of the proinflammatory chemokines *Cxcl2* and *Ccl9* (**Fig. 3e, h**), as well as marked increases in the immune response genes *H2-Aa*, *H2-Ab1*, *H2-Eb1*, and *Cd74* (**Fig. 3e**). For example, elevated levels of *Cd74*, a class II histocompatibility antigen gamma chain, have been previously observed on monocytes within atherosclerotic plaques ^38^. The relative expression levels of each of these genes were lower in the *Tet2+/+*;CD45.2+ wild-type monocytes than in *Tet2+/-*;CD45.1+ mutant monocytes, and eltanexor treatment reduced the expression of each of these genes in *Tet2+/-*;CD45.1+ monocytes (**Fig.3e**,**h**), indicating an anti-inflammatory effect of eltanexor in addition to its profound effect on reducing the numbers of the *Tet2+/-* ;CD45.1+ monocytes (**Fig. 3b top**, **3c**).

We also analyzed cluster 5, which we classified as M2 macrophages (**Fig. 3b**, **3f, Extended Data Fig. 4**). In *Tet2+/-*;CD45.1+ M2 macrophages, we found aberrant upregulation of the proinflammatory chemokines *Ccl4* and *Ccl7,* both potent monocytes^39,40^ chemoattractants (**Fig.3f**), as well as the chemokine receptor *Ccr2* (**Fig.3i**) compared to *Tet2+/+*;CD45.2+ M2 macrophages. *Klf4* also rose in the *Tet2+/-*;CD45.1+ M2 macrophages (**Fig.3f, Extended Data Fig. 6**), mirroring the pattern observed in *Tet2+/-*;CD45.1+ M1 macrophages. Importantly, eltanexor treatment reduced the expression of these proinflammatory genes in *Tet2+/-*;CD45.1+ M2 macrophages, similar to the findings for M1 macrophages (Cluster 2) and monocytes (Cluster 8) (**Fig.3f, i, Extended Data Fig.6**), Taken together, these data indicate that eltanexor treatment reduced both the total numbers of *Tet2+/-*;CD45.1+ monocytes and the proinflammatory cytokine and chemokine expression by *Tet2+/-*;CD45.1+ monocytes and macrophages within the aortic arch, which may explain the reduced formation of atherosclerotic plaques observed in eltanexor-treated Group 1 mice transplanted with *Tet2+/-*;CD45.1+ mutant bone marrow cells (**Fig. 1**).

### Mice receiving *Tet2+/-*;CD45.1*+* mutant hematopoietic cells exhibit a hyperinflammatory profile in the cells of the microenvironment of atherosclerotic plaques, which is reduced by eltanexor treatment

Vascular smooth muscle cells, endothelial cells and fibroblasts contribute crucially to atherogenesis ^41–43^. To elucidate the impact of *Tet2*-mutant hematopoietic cells on the plaque microenvironment, we determined the gene expression of CD45-negative, non-hematopoietic cells in Group 1 mice injected with a mixture of *Tet2+/-*;CD45.1*+* and *Tet2+/+*;CD45.2+ bone marrow cells (**Fig. 1a**), and compared to them to those in Group 2 control mice, which received a mixture of *Tet2+/+*;CD45.1*+* and *Tet2+/+*;CD45.2+ bone marrow cells (**Fig. 4).**

**Figure 4.**
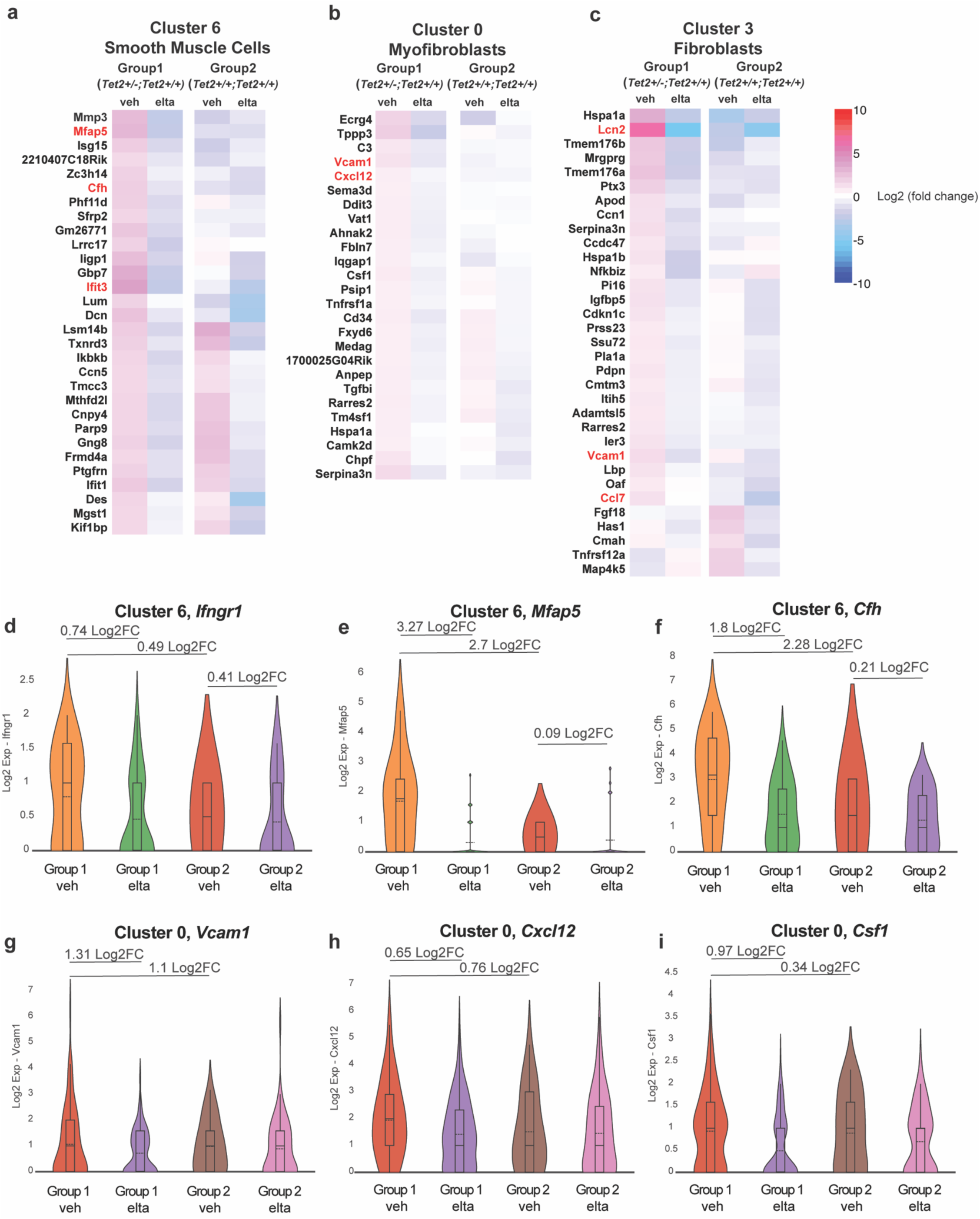
Mice receiving *Tet2+/-*;CD45.1*+* mutant hematopoietic cells exhibit a hyperinflammatory profile in the microenvironment of atherosclerotic plaques, which is reduced by eltanexor treatment. Mice were treated as indicated in Figure 1a**. a-c**, Heatmaps of differentially expressed genes in non-hematopoietic cells from eltanexor treated and vehicle control mice injected with *Tet2+/-*;CD45.1+ bone marrow cells (Group 1) relative to non-hematopoietic cells from mice injected with *Tet2+/+*;CD45.1+ bone marrow cells (Group 2) for (a) smooth muscle cells (cluster 6), (b) myofibroblasts (Cluster 0) and (c) fibroblasts (Cluster 3). **d-i**, Violin plots of scRNA-seq data from non-hematopoietic cell populations from eltanexor-treated and vehicle-treated control mice injected with *Tet2+/-*;CD45.1+ bone marrow cells (Group 1) relative to non-hematopoietic cells from mice injected with *Tet2+/+*;CD45.1+ bone marrow cells (Group 2) for (d) *Ifngr1* (e) *Mfap5* (f) *Cfh* in smooth muscle cells (cluster 6), and (g) *Vcam1*(h) *Cxcl12* and (i) *Csf1* in fibroblasts (cluster 0).

In the vascular smooth muscle cells from the aortic wall (Cluster 6), which we identified based on the expression of *Acta2*, *Cnn1*, and *Myh11* ^44^ (**Extended Data Fig. 7a-c**), we detected 15 genes with upregulated RNA expression in vehicle-treated Group 1 mice injected with *Tet2+/-* ;CD45.1*+* cells, but not in vehicle-treated Group 2 mice injected with *Tet2+/+*;CD45.1+ cells (**Fig. 4a**). Importantly, eltanexor treatment reduced the expression of each of these 15 genes (**Fig. 4a)**, which might contribute to the smaller atherosclerotic plaques in eltanexor-treated mice (see **Fig. 1b,c**). Among these 15 genes, the gene encoding interferon-induced protein with tetratricopeptide repeats 3 (*Ifit3*) stands out as the most differentially expressed gene in vehicle-treated Group 1 mice versus vehicle-treated Group 2 mice (**Fig. 4a**). *Ifit3* encodes an interferon-induced protein whose high expression levels occur in patients with early onset atherosclerosis ^45^. We additionally found increased RNA that encodes interferon-gamma receptor 1 (*Ifngr1*) in smooth muscle cells of vehicle-treated Group 1 mice compared to vehicle-treated Group 2 mice (**Fig. 4d**). These findings suggest that high *Ifit3* and *Ifngr1* expression levels in smooth muscle cells of vehicle-treated mice injected with *Tet2+/-*;CD45.1+ cells (Group1) result from the increased inflammatory signaling by *Tet2*+/- mutant cells. Importantly, treatment with eltanexor markedly reduced *Ifit3* expression as well as *Ifngr1* expression in Group 1 mice injected with *Tet2+/-*;CD45.1+ bone marrow cells, indicating that the anti-inflammatory effects of eltanexor not only influence the mutant blood cells but also extend to host smooth muscle cells (**Fig. 4a, d**). *Mfap5,* a gene upregulated in obesity-related inflammation^46^, and Complement factor H (Cfh), a gene whose increased expression exacerbates atherosclerosis ^47^, also exhibited increased expression in vascular smooth muscle cells of mice injected with *Tet2+/-*;CD45.1*+* bone marrow cells and decreased expression with eltanexor treatment (**Fig. 4e, f**).

We classified cells in Cluster 9 as vascular endothelial cells, based on the expression of *Pecam1* (CD31), *Vwf*, *Flt1*, and *Cdh5* (**Extended Data Fig. 7d-g**)^48^. While we were not able to compare endothelial cells of vehicle-treated mice of Group 1 with those of vehicle-treated mice of Group 2, due to relatively low numbers of endothelial cells recovered from the latter, we did observe expression of proinflammatory chemokines and chemokine receptors including *Ccl2* and *Cxcl12* in endothelial cells of mice injected with *Tet2+/-*;CD45.1+ bone marrow cells, which eltanexor treatment substantially reduced (**Extended Data Fig. 8a, b**). Moreover, eltanexor reduced expression of the immune response genes *H2-Aa, H2-Ab1*, and *CD74* in these cells (**Extended Data Fig. 8a,c**), similar to our findings in *Tet2+/-*;CD45.1+ monocytes (**Fig. 3e**). Cardiomyocytes from vehicle-treated Group1 mice injected with *Tet2+/-*;CD45.1+ bone marrow, had increased expression of *Fos,* a gene that is critical for foam cell formation ^49^, and *Igfbp5* which has been previously found to be increased in atherosclerotic plaques ^50^. The expression of both genes fell substantially with eltanexor treatment (**Extended Data Fig. 8d**). Next, we focused on Clusters 0 and 3, which we classified as myofibroblasts and fibroblasts, respectively, based on the expression levels of *Acta2*, *Dcn*, *Col1a1,* and *Pdgfra* (**Extended Data Fig. 7a,h-j)**^48^. Fibroblasts of mice injected with *Tet2+/-*;CD45.1+ bone marrow cells had increased transcripts that encode *Ccl7*, a chemokine associated with atherosclerosis^51^, that fell with eltanexor treatment (**Fig. 4c**). Myofibroblasts of these mice had augmented expression of the genes encoding stromal cell-derived factor 1 (*Cxcl12*), macrophage colony-stimulating factor 1 (*Csf1*), and vascular cell adhesion molecule 1 (*Vcam1*), all of which eltanexor treatment reduced (**Fig. 4b,g-i**). *CSF1* and *CXCL12* both likely participate causally coronary artery disease (CAD) in humans ^52^. VCAM1 critically contributes to atherosclerosis by mediating monocyte infiltration into the intima and VCAM-1 inhibition effectively blocked monocyte infiltration and consequently atherosclerotic lesion formation ^53^.

Taken together, our results indicate that smooth muscle cells, endothelial cells, and fibroblasts of mice injected with *Tet2+/-*;CD45.1*+* bone marrow cells and treated with vehicle exhibit increased expression of inflammatory mediators which promote monocyte infiltration into the intima to accelerate atherosclerotic plaque formation. The concomitant reduced expression in each of these genes due to eltanexor treatment further supports this drug as a candidate to reducing both inflammation and atherosclerotic plaque formation.

### *Atf3* occupies the largest super-enhancer in wild-type macrophages, and ATF3 DNA binding is reduced in *Tet2*-mutant bone marrow-derived macrophages

The mechanisms underlying the enhanced pro-inflammatory palette of *Tet2*-mutant macrophages and whether dysregulation of cell-type-specific enhancers due to Tet2 loss results in increased inflammation are poorly understood. Previous studies have found that *TET2*-deficiency is associated with hypermethylated enhancers in CHIP^4^ and that deletion of *Tet2* in murine embryonic stem cells results in changes in transcription factor binding within regions of open chromatin associated with cell-type–specific enhancers ^54^. These studies provide examples of how *Tet2* loss-of-function can affect gene expression patterns and drive disease development. To investigate the enhanced proinflammatory profile of *Tet2*-mutant macrophages and evaluate the role of *Tet2* loss in mediating enhancer dysregulation, we first performed enhancer profiling based on ChIP-seq for H3K27ac in bone marrow-derived macrophages (BMDM) from *Tet2+/+, Tet2+/-* or *Tet2-/-* mice (**Fig. 5a**). *Tet2+/+* BMDMs displayed 18 transcription factors (TFs) that are regulated by large *cis*-regulatory elements called stretch- or super-enhancers (SE) that drive high levels of gene expression from target promoters (**Fig. 5b**) ^55,56^. Interestingly, the number of transcription factors associated with SEs increased to 21 in *Tet2+/-* BMDMs and 30 in *Tet2-/-* BMDMs (**Fig. 5b**). *Atf3* was associated with the largest SE in *Tet2+/+* wild-type BMDMs, and the levels of H3K27ac within this SE, as well as its mRNA expression levels fell in *Tet2+/-* and *Tet2-/-* BMDMs (**Fig. 5c-e**). ATF3, which inhibits the macrophage inflammatory response, is a transcription factor that can either bind as a homodimer, where it is reported to act as a repressor, or as a heterodimer with CEBPA or other CREB family members including ATF4, JUN, JUNB or JUND, where it can activate transcription ^29,57^. Notably, we found that the H3K27ac signal of *Cebpa*, a transcription factor associated with AML when downregulated and shown to counteract inflammation when upregulated ^58^, also decreased in *Tet2+/-* and *Tet2-/-* BMDMs. (**Fig. 5b,c**). *Cebpa* mRNA expression levels were high in wild-type *Tet2+/+* BMDMs, and significantly reduced in *Tet2+/-* and *Tet2-/-* BMDMs, consistent with the H3K27ac profile we observed in each of the three genotypes (**Fig. 5f**). By contrast, *Atf4* and *Junb* mRNA levels increased in *Tet2+/-* and *Tet2-/-* BMDMs, whereas c-*Jun* and *Jund* mRNA expression levels did not show a consistent pattern across the three genotypes (**Fig. 5g-j**).

**Figure 5.**
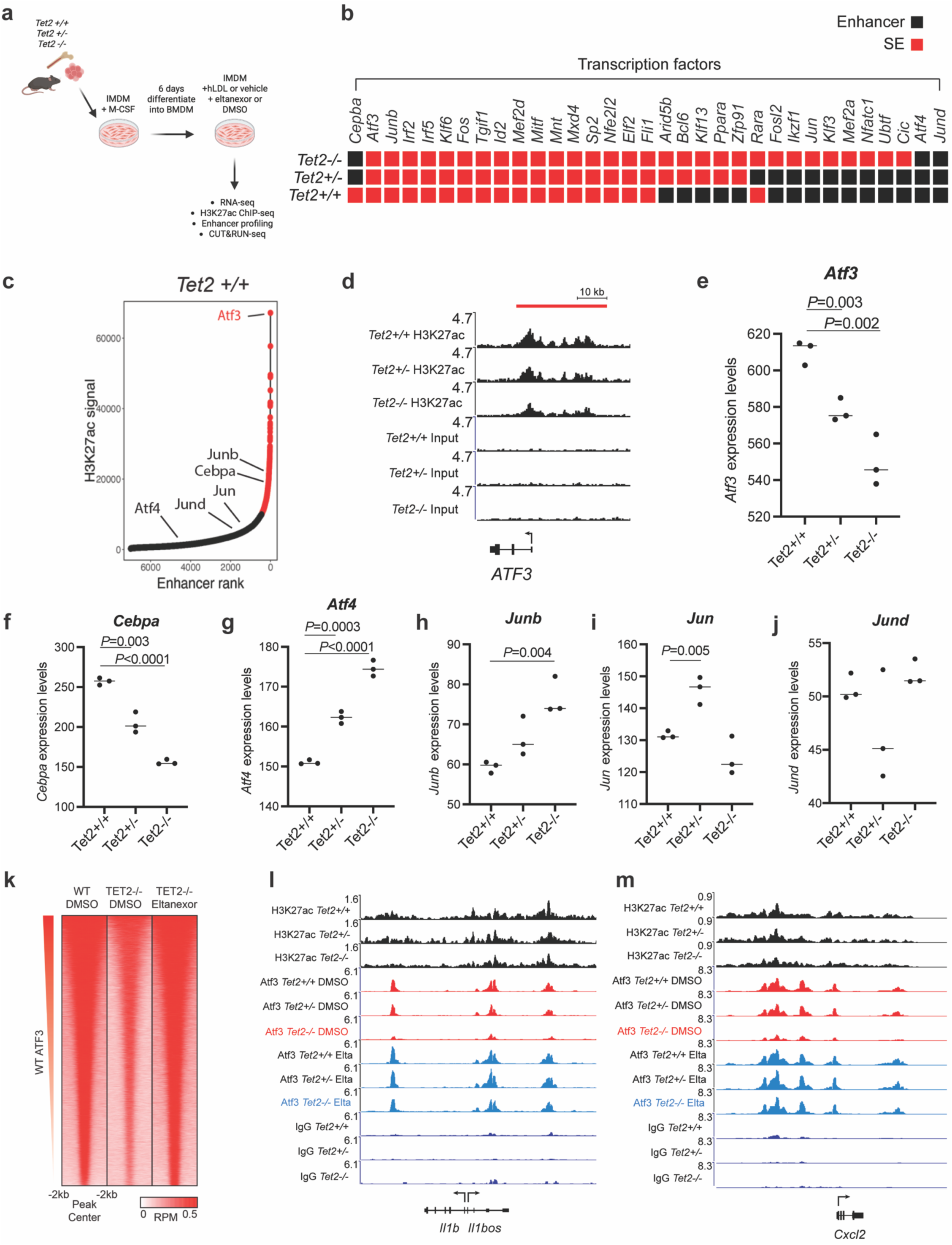
*Atf3* occupies the largest super-enhancer in wild-type macrophages, and ATF3 DNA binding is reduced in *Tet2*-mutant bone marrow-derived macrophages. **a,** Macrophages were obtained from the bone marrow of *Tet2+/+*, *Tet2+/-* or *Tet2-/-* mice and cultured for 6 days in IMDM plus M-CSF followed by incubation for 24 hours with hLDL (200 mg/dl) and either eltanexor (100 nM) or vehicle control. Bone marrow-derived macrophages (BMDM) were then harvested and processed for ChIP-seq, CUT&RUN and RNA-seq experiments. **b**, Enhancer profiling of BMDM from *Tet2+/+*, *Tet2+/-* or *Tet2-/-* mice revealed a conserved set of SEs associated with genes encoding transcription factors (n=31). The enhancers are highlighted in red in mouse genotypes when they are large enough to qualify as SEs, and in black when they do not based on the H3K27Ac signal. **c**, Ranking of enhancers by H3K27ac signal associated with genes in BMDM from *Tet2+/+* mice. *Atf3* was associated with the largest SE in *Tet2+/+* BMDMs. Atf3 binding partners Atf4, Jun, Junb, Jund, and Cebpa are also indicated. **d**, Normalized ChIP-seq alignment tracks for H3K27ac at the *Atf3*-associated SE in BMDM from *Tet2+/+, Tet2+/-* or *Tet2-/-* mice. ChIP-seq read densities (y-axis) were normalized to reads per million reads sequenced from each sample. Red bar indicates the location of SEs. **e-j**, Gene expression levels by RNAseq of *Atf3* (e), *Cebpa* (f) *Atf4* (g), *Junb* (h) *Jun* (i) *Jund* (j) in BMDM from *Tet2+/+*, *Tet2+/-* or *Tet2-/-* mice. **k**, Genome-wide occupancy for ATF3 from control *Tet2+/+* and *Tet2-/-* BMDM and from eltanexor-treated *Tet2-/-* BMDM determined by CUT&RUN. Genomic regions (rows) were defined as those enriched in sequencing reads for at least one condition and are ranked by the ATF3 signal across the region. **l-m**, CUT&RUN coverage tracks for ATF3 and IgG (control) in DMSO control- or Eltanexor (Elta)-treated BMDM from *Tet2+/+, Tet2+/-* or *Tet2-/-* mice, overlaid with H3K27ac ChIP-seq at the *Il1b* locus (l) and the *Cxcl12* locus (m).

To identify regions of sequence-specific genomic occupancy of ATF3, we next performed CUT&RUN analysis ^59^. Consistent with its decrease in RNA expression, ATF3 binding was reduced across the genome in *Tet2-/-* BMDMs as compared to *Tet2+/+* BMDMs. Interestingly, eltanexor treatment restored this loss of binding (**Fig. 5k**). As Atf3 is a major inhibitor of inflammatory genes, including those that promote atherosclerosis ^29,30,60–62^, we examined ATF3 binding at genes encoding pro-inflammatory cytokines that were upregulated in aortic root macrophages from mice injected with *Tet2*-mutant cells (see **Fig. 3**). Our investigation of wild-type BMDMs revealed that ATF3 bound the enhancers of *Il*-*1β* (**Fig. 5l**), *Cxcl2* (**Fig. 5m**) *Il*-*6*, *Il*-*12rb*, *Cxcl10*, *Ccl4*, *Ccl9*, *Ccl12*, and *Tnfaip2* (**Extended Data Fig. 9**). Interestingly, this binding was significantly diminished in *Tet2*-mutant BMDMs, consistent with a mechanism in which ATF3 represses the expression of these inflammatory genes in wild-type macrophages but not in *Tet2*-mutant macrophages. Strikingly, eltanexor treatment of *Tet2*-mutant BMDMs completely restored ATF3 binding levels to those of wild-type BMDMs, consistent with increased repression of inflammatory gene expression (**Fig 5k, L, M, Extended Data Fig 9).**

### Reduced binding of ATF3 to the enhancers of pro-inflammatory mediators occurs in *Tet2-*mutant BMDM without evidence of aberrant methylation of CpGs

TET2 is a cyclooxygenase that catalyzes the oxidation of 5-mC to 5-hmC, the first step in removing methylation from key CpGs within the enhancers of expressed genes ^63^. The preferred binding sites, i.e. motif occurrences, of ATF3 bear a CpG, which may be methylated in certain contexts. Other CREB/ATF family members^57^, namely ATF4^64^, show differential preference for unmethylated and methylated DNA. We therefore hypothesized that some or all ATF3 motif occurrences within inflammatory gene enhancers are differentially methylated in *Tet2* wild-type versus mutant cells and thus affect binding of ATF3 to the enhancers of key pro-inflammatory cytokine genes. To determine whether the reduced binding of Atf3 to the enhancers of inflammatory cytokine genes in *Tet2*-mutant macrophages results from aberrant CpG methylation, we performed whole-genome-wide bisulfite sequencing (WGBS) in *Tet2+/+*, *Tet2+/-*, and *Tet2-/-* BMDMs. Overall, and consistent with the results of other investigators ^4^, we found that *Tet2-/-* macrophages exhibited largely unchanged global methylation, but where changes were induced, methylation tended to increase over the wild-type Tet2+/+ macrophages (**Fig. 6a)**. To examine the possibility that differential methylation affected the CpGs within ATF3- binding sites at the enhancers of inflammatory genes, we overlaid the WGBS tracks for *Tet2+/+*, *Tet2+/-* and *Tet2-/-* BMDMs with H3K27ac CHIP-seq tracks as well as CUT&RUN-seq tracks for Atf3 at the gene loci of *Il-1b, Il12rb2, Ccl4, Ccl9, Cxcl10,* and *Tnfaip2* (**Fig. 6b-g**). As shown in **Fig. 6b-g**, we did not identify consistently differentially methylated CpGs at ATF3-binding sites within the enhancers of these genes, indicating that differential methylation of these ATF3 binding sites was not the prevailing cause for altered ATF3 binding.

**Figure 6.**
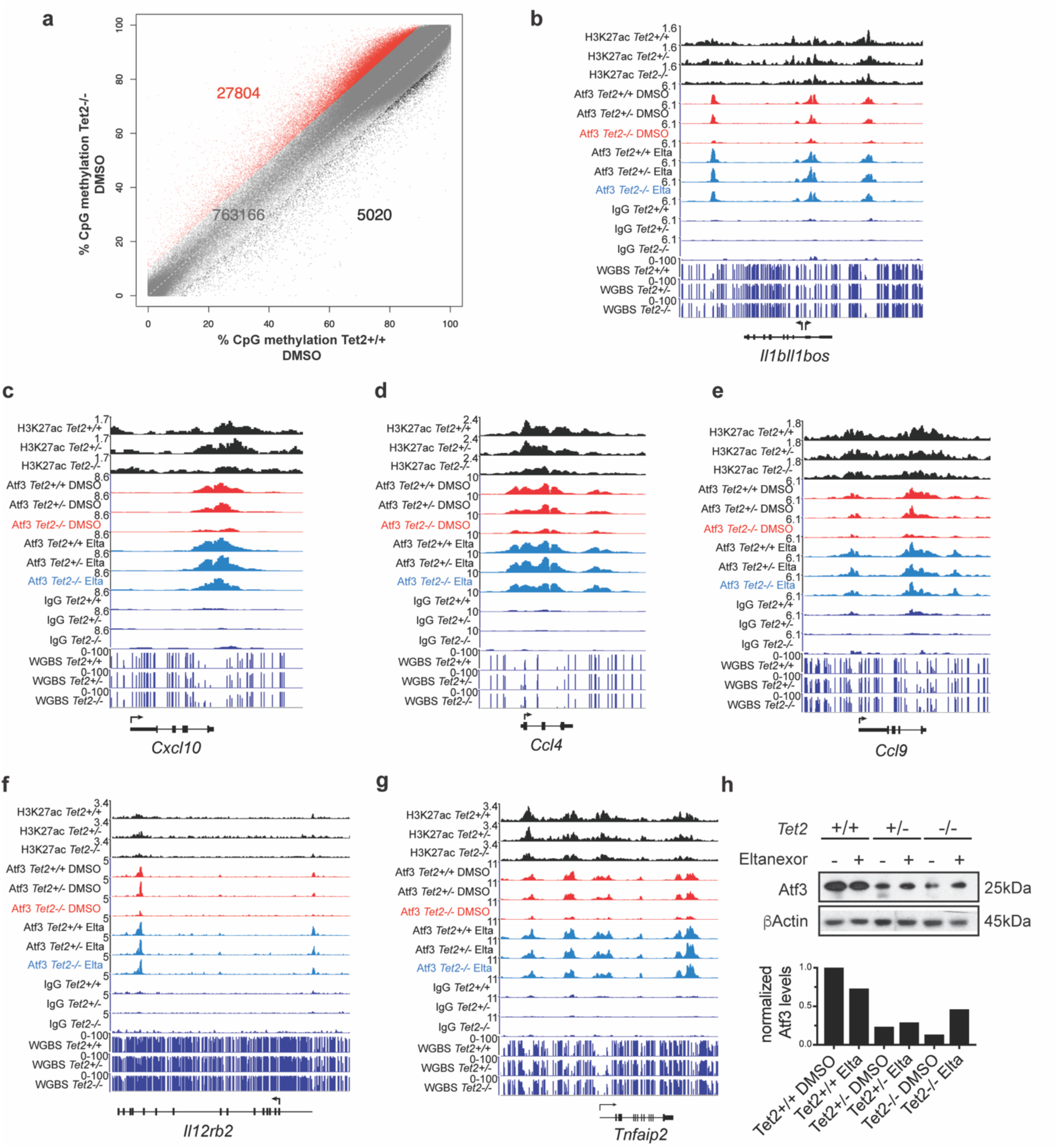
Reduced expression of ATF3, occurs in *Tet2-*mutant BMDM, without evidence of aberrant methylation of CpGs within the enhancers of pro-inflammatory mediators. **a,** Differentially methylated regions (DMRs) from 50-CpG-long regions of the genome (> 10% methylation difference) between *Tet2-/-* and *Tet2+/+* BMDMs . **b-g**, Whole genome bisulfite sequencing (WGBS) tracks for Tet2+/+, Tet2+/- and Tet2-/- BMDMs are shown in the bottom three rows for the *IL1b* locus (b), the *Cxcl10* locus (c), the *Ccl4* locus (d), the *Ccl9* locus (e), the *IL12rb2* locus (f), and the *Tnfaip2* locus (g). These tracks are overlaid with the CUT&RUN coverage tracks for ATF3 and IgG (control) and for H3K27ac ChIP-seq. **h**, (Top) Western blot of BMDM from *Tet2+/+, Tet2+/-* or *Tet2-/-* mice with or without eltanexor treatment using antibodies specific for ATF3, and beta-actin. (Bottom) Normalized Atf3 levels from densitometric quantification.

Next we examined the protein levels of ATF3 as a possible cause of reduced binding of ATF3 to the regulatory regions of inflammatory genes in *Tet2*-mutant cells. Correlating with the reduced mRNA expression levels in *Tet2*-deficient BMDM (see **Fig. 5e**), the protein levels of ATF3 fell with *Tet2* loss (**Fig. 6h**), offering an explanation for reduced ATF3 binding in these cells. Interestingly, eltanexor treatment partially restored ATF3 protein levels in *Tet2* mutant BMDMs, explaining in part the restored binding of ATF3 to its binding sites in the CUT&RUN assays in eltanexor-treated *Tet2*-deficient BMDMs (**Fig. 5 l-m, Fig. 6b-g, Extended Data Fig. 9a-g).**

## Discussion

Our studies in murine models of *Tet2*-deficient CHIP show that eltanexor, an inhibitor of the nuclear export protein XPO1, markedly reduced atherosclerotic plaque formation in mice receiving *Tet2*-mutant bone marrow cells, with a two-fold decrease in total lesion volume as compared to vehicle-treated mice, and a 1.4-fold reduction in the percentage of CD68+ macrophages into the atheroma (**Fig. 1b-d**). Thus, eltanexor joins the small group of targeted agents that have been tested for their ability to reduce the risk and extent of atherosclerosis in experimental models of *TET2*-mutant CHIP, including IL1β-blockers such as canakinumab or anakinra ^10,17,65^, IL6R-blocking antibodies ^11^, inhibitors of the NLRP3 inflammasome, such as MCC950 ^18^, and colchicine, a microtubule inhibitor with anti-inflammatory properties ^19^. Eltanexor is a second-generation XPO1 inhibitor similar to Selinexor but better tolerated due to its inability to cross the blood-brain barrier^66^. It has been evaluated in a phase 1/2 clinical trial in patients with hematologic and solid tumors^67^ and we previously showed that eltanexor mediates selective killing of *Tet2*-mutant HSPCs without inducing DNA damage^21^. Both eltanexor and selinexor block the nuclear-cytoplasmic transport of specific proteins, as well as mRNAs encoding proteins involved in inflammation, self-renewal, and cell proliferation ^25^.

Our single-cell CITE-seq analysis of competitively transplanted mice suggests that the reduction in atherosclerotic plaque formation in eltanexor-treated mice stems not only from a decrease of CD68+ *Tet2*-mutant macrophages within the plaque (**Fig. 1d,e**) but also from a significant dampening of proinflammatory cytokine expression in both peripheral macrophages and bone marrow monocyte progenitors (**Fig. 2 and 3**). For instance, we found augmented expression of numerous inflammatory mediators, including *Il1b*, *Nfkb1*, *Ccl2*, *Ccl7*, *Ccl9*, *Ccl12*, *Cxcl10*, *CD74*, and *Klf4* in *Tet2*-mutant macrophages and monocytes *in vivo* (**Fig. 3d-I, Extended Data Fig. 6**), several of which were reported increased by other groups in experimental models of *Tet2* -mutant CHIP ^2,7,13,23^, as well as in human atherosclerotic plaques^68^. Eltanexor treatment reduced expression of each of these proinflammatory cytokines, chemokines, and immune response genes to the levels expressed by normal macrophages (**Fig. 3d-I, Extended Data Fig. 6**).

Our findings regarding macrophage progenitors in the bone marrow showed increased expression levels of inflammatory genes including *Aif1* (**Fig. 2d, g**), which likely reflects activation by proinflammatory cytokines secreted by surrounding mutant monocytes and macrophages ^28^. Moreover, *Atf3*, a key transcriptional regulator that inhibits the inflammatory response in normal macrophages ^61,62,69^, rose with eltanexor treatment in *Tet2*-mutant monocyte macrophage progenitors (**Fig. 2h**). Thus, eltanexor not only reduces the production of proinflammatory cytokines by *Tet2*-mutant monocytes and macrophages but also augments the expression of genes like ATF3 that normally act to attenuate the proinflammatory response ^30^.

Eltanexor significantly reduced the increased expression of the self-renewal gene *Ltb* in *Tet2*-mutant HSCs ^23,24^ (**Fig. 2d-f**), which may contribute to the ability of eltanexor to mitigate the clonal expansion of *Tet2*-mutant HSCs (**Figure 2b,c**). Indeed, we observed a slight decline in the relative numbers of *Tet2*-mutant HSCs and a substantial decline in *Tet2*-mutant CMPs in the bone marrow with eltanexor treatment (**Fig. 2b, c**). Further studies over a longer time frame will be needed to demonstrate whether eltanexor can ultimately delay or prevent the malignant transformation of *Tet2*-mutant myeloid progenitors. Nevertheless, the simultaneous suppression of *Ltb* and *Aif1* and the rise of *Atf3* in *Tet2+/-* HSPCs post-treatment with eltanexor indicate both a restoration of more normal hematopoietic cell homeostasis and a reduction in pro-inflammatory signaling pathways that promote atherosclerosis, ultimately disrupting the inflammatory feedback loop in *Tet2*-deficient CHIP.

The single-cell RNA studies also probed CD45-negative intrinsic arterial cells within the atherosclerotic plaques. Mice injected with *Tet2+/-;*CD45.1+ mutant hematopoietic cells (Group 1) exhibited a hyperinflammatory slant of non-leukocyte plaque cells (**Fig. 4**), compared with control mice injected with *Tet2+/+;*CD45.1+ normal bone marrow cells (Group 2). For example, vascular smooth muscle cells from the aortic arch of vehicle-treated Group 1 mice showed increased expression of at least 15 genes compared to vehicle-treated Group 2 control mice, including *Ifit3*, *Mfap5*, and *Cfh*, and each of these transcripts was downregulated following eltanexor treatment (**Fig. 4a, e-f**). Vascular endothelial cells of Group 1 mice also exhibited elevated expression of pro-inflammatory chemokines such as *Ccl2* and *Cxcl12*, as well as the immune response genes *H2-Aa* and *H2-Ab1*, and the increased expression of each fell substantially with eltanexor (**Extended Data Fig. 8 a-c**). In the aortic plaque fibroblasts of Group 1 mice injected with *Tet2*-mutant bone marrow cells, eltanexor reduced the elevated expression of genes associated with inflammation and monocyte infiltration, including *Csf1*, *Cxcl12*, and *Vcam1* (**Fig. 4b,c,g-i**). Smooth muscle cells, endothelial cells, and fibroblasts contribute critically to atherosclerotic plaque formation ^41–43^. For instance, vascular smooth muscle cells proliferate and migrate into the atherosclerotic plaque to contribute with macrophages to foam cell formation ^70^, and endothelial cells recruit inflammatory cells and modify lipoprotein particles ^71–73^. Moreover, fibroblasts also participate in the inflammatory response, and with smooth muscle cells, in extracellular matrix production ^65,74^. Our single-cell CITE-seq study in a competitive repopulation model of *Tet2*-deficient CHIP clearly demonstrates that the proximity of *Tet2*-mutant macrophages changes the gene expression profile of nonhematopoietic cells in a paracrine manner to accelerate atherosclerotic plaque formation. Moreover, our findings highlight that the anti-inflammatory effects of eltanexor extend beyond *Tet2*-mutant macrophages and hematopoietic cells, targeting the microenvironment to reduce inflammation and atherosclerotic plaque formation further.

Several studies have reported altered transcription factor recruitment to specific enhancers in *Tet2*-mutant embryonic stem cells and myeloid hematopoietic cells, leading to a better understanding of how Tet2 loss affects gene expression and promotes disease development ^4,75^. However, the underlying mechanisms causing the hyperinflammatory profile of *Tet2-*mutant macrophages remain incompletely understood. Our enhancer profiling studies in wild-type and *Tet2*-mutant bone marrow-derived macrophages revealed that *Tet2* loss-of-function mutations lead to an increased number of genes encoding transcription factors that associate with stretch or super-enhancers in *Tet2*-mutant macrophages, indicating heightened transcriptional activity (**Fig. 5b**). Strikingly, *Atf3* emerged as the transcription factor gene linked to the largest super-enhancer in wild-type macrophages (**Fig. 5c**). *Tet2*-loss substantially reduced *Atf3*-associated H3K27ac signal, mRNA expression and protein expression (**Fig. 5d, e, Fig. 6h**), highlighting ATF3 as a potentially central, *Tet2*-sensitive regulator in CHIP. CUT&RUN analyses further showed a significant reduction in ATF3 binding to the enhancers of numerous genes encoding inflammatory cytokines and chemokines that are overexpressed in *Tet2-/-* macrophages (**Fig. 5e-g, Extended Data Fig. 9**), suggesting that ATF3 functions as a transcriptional repressor of these pro-inflammatory genes. Importantly, eltanexor treatment restored ATF3 binding levels at these enhancers in *Tet2*-mutant macrophages, providing a mechanism for the eltanexor-mediated reduction of expression of inflammatory genes that are not themselves XPO1 cargos but rather appear to be transcriptionally regulated by ATF3.

To understand the reduced binding of ATF3 to enhancers of inflammatory genes in *Tet2*-mutant macrophages, we performed bisulfite sequencing and confirmed a trend toward increased methylation as previously reported by others ^54,75^ (**Fig. 6a**). However, we did not identify consistent patterns of differential CpG methylation at ATF3 binding sites within the enhancers of upregulated pro-inflammatory cytokines, suggesting that impaired ATF3 recruitment does not result from methylation changes of these regions (**Fig. 6b-g**). While reduced *Atf3* mRNA and protein expression in *Tet2*-mutant macrophages likely contributes to decreased enhancer occupancy (**Fig. 5e**, **Fig. 6h**), additional factors may affect ATF3 binding activity in *Tet2*-deficient cells. ATF3 is known to heterodimerize with JUN, JUNB, JUND, and/or ATF4, to form context-specific transcriptional complexes that can either repress or activate the transcription of inflammatory genes ^29,30,57^. Further studies are needed to assess whether disruption of these interactions or altered expression of ATF3 binding partners contribute to impaired ATF3 function in *Tet2*-deficient cells.

In summary, our findings identify ATF3 as a *Tet2*-sensitive master regulator of inflammation in CHIP and demonstrate the therapeutic potential of eltanexor as a targeted agent that disrupts the inflammatory feedback loop in *Tet2*-deficient macrophages by restoring ATF3 binding to the enhancers of inflammatory genes, thereby decreasing their expression. This action significantly reduces atherosclerosis through the attenuation of pro-inflammatory signaling in macrophages of eltanexor-treated mice, and the modulation of non-hematopoietic cell responses within the plaque microenvironment. Eltanexor acts through a multifaceted mechanism, simultaneously reducing the expression of inflammatory cytokines, chemokines, transcription factors, and self-renewal genes in *Tet2*-mutant hematopoietic cells, ultimately rebalancing cellular homeostasis and reducing atherosclerosis. Taken together, these actions comprehensively normalize the disease landscape of *Tet2*-mutant CHIP. These preclinical results justify the initiation of clinical trials to determine whether eltanexor treatment reduces the progression of atherosclerosis and the risk of heart attack or stroke in patients with *TET2*-mutant CHIP. Furthermore, elucidating the mechanisms of altered ATF3 function in *Tet2*-mutant CHIP will require additional studies to advance our understanding of *TET2*-related mechanisms and inform future therapeutic strategies.

## Acknowledgments

This work was supported by NIH grants R35CA210064 (A.T.L.), The G. Harold and Leila Y. Mathers Charitable Foundation (A.T.L., N.P.), Karyopharm (A.T.L., N.P.) and the St. Jude Children’s Research Hospital Transcription Collaborative (A.T.L., B.J.A.). N.P was supported by the CCEH-NIDDK P&F Award and the EvansMDS Young Investigator Award. A.V. received the Harold M. English Fellowship Fund from Harvard Medical School (Boston, USA). B.J.A. was supported by the American Lebanese Syrian Associated Charities (ALSAC). M.W.Z. was supported by grants from Alex’s Lemonade Stand Foundation, Charles A. King Trust, and Claudia Adams Barr Foundation. D.N. reports stock ownership in Madrigal Pharmaceuticals. P. L. receives funding support from the National Heart, Lung, and Blood Institute (R01HL170000, 1R01HL163099-01, R01AG063839, R01HL151627, R01HL157073, R01HL166538), and the RRM Charitable Fund. We thank Ben Ebert for his intellectual guidance on the mouse study. We also thank the Neurobiology Imaging Facility of Harvard Medical School for the access to the Olympus slide scanner VS-120. We also thank John Easton and Heather Mulder from St. Jude, for assistance with whole genome bisulfite sequencing experiments.

## Methods

### Mice

*Tet2*-deficient mice (B6(Cg)-*Tet2^tm1.2Rao^*/J (strain no. 023359; CD45.2) were initially crossed with Pep Boy mice (strain no. 002014; CD45.1) from The Jackson Laboratory (ME, USA) to generate Tet2-deficient mice (Tet2 +/-) in the CD45.1 background. *Ldlr*^−/−^ mice (strain no. 002207; CD45.2) and C57BL/6J (strain no. 000664; CD45.2) were also obtained from The Jackson Laboratory (ME, USA). All animal experiments were conducted in accordance with the Institutional Animal Care and Use Committee of the Dana Farber Cancer Institute.

### Bone marrow transplantation and drug administration and high cholesterol diet

Whole bone marrow cells were isolated from femurs, hips, and tibias from 8-12 weeks old donor mice, (Tet2 +/+, CD45.2 or Tet2 +/-, CD45.1) by centrifugation at 8,000*g* for 1 min. Female recipient mice aged 6 weeks (*Ldlr*^−/−^ mice) were lethally irradiated with split doses 475 cGy (4 hours apart for a total dose of 950 cGy) using X-Rad 225Cx Irradiator System (Precision X-Ray, CT). Within 1 hour of the last radiation dose, bone marrow cells from the donor mice (1 million cells from Tet2 +/- or Te2 +/+, CD45.1 mixed with 1 million cells from Tet2 +/+, CD45.2) were injected intravenously. Eight weeks after engraftment, mice were randomized into treatment cohorts of vehicle control (0.5% methylcellulose with 1% Tween 80 in water) or eltanexor (5 mg/kg on a 5 days ON 2 days OFF schedule, Karyopharm Therapeutics, MA) and treated for 12 weeks by oral gavage. All mice were fed a high-cholesterol diet (Envigo, TD.88137 diet, IN) and water *ad libitum*. Body weight measurements were recorded twice a week over the course of 12 weeks.

### Bone Marrow Derived Macrophages

Whole bone marrow was isolated from long bones, hips, and vertebrae of 8-12 week old mice by crushing and sequential passage through 70 μm cell strainers (Corning Cat. No. 352350 and 352340). Red cell lysis with ACK Lysing Buffer (Gibco Cat. No. 10492-01) was performed and bone marrow was cultured by creating a single-cell suspension of whole bone marrow in Iscove’s Modification of DMEM (IMDM) (Corning Cat. No. 10016CV) supplemented with 10% fetal bovine serum (FBS) (Omega Scientific Cat. No. FB- 11), 10 ng/mL recombinant mouse macrophage colony-stimulating factor (M- CSF, Miltenyi Biotec Cat. No. 130-101-706), and 1% penicillin/streptomycin/glutamine (PSG) (Gibco Cat. No.10378-016) in 30 mL total volume. After 3 days, each dish was supplemented with 15 mL of the above media. On day 6, bone marrow-derived macrophages (BMDM) were stimulated with media containing 200 mg/dL LDL and either 50 mm eltanexor or vehicle. After 24 hours, macrophages were harvested with a cell lifter and used for ChIP-seq, CUT&RUN, RNA-seq, or Western Blot experiments.

### Atherosclerosis lesion analysis and metabolic profiling

Heart tissue was isolated from chimeric mice after 12 weeks of treatment and diet, embedded in Optimal Cutting Temperature compound (OCT) and frozen. Aortic roots were serially sectioned at 6mm thickness. Six sections per mouse were stained with Oil red for total lesion area quantification. Data were collected using an Olympus VS120 Slide Scanner at 20x magnification. Quantification was performed using ImageJ. Total plasma cholesterol was measured using a kit from Wako Diagnostics (Cat.No 10752-436, VWR).

### Immunohistochemistry staining

Six sections per mouse were stained with anti-CD68 (1:400 dilution, Cat. no. 137001, Biolegend). The sections were incubated with primary antibodies overnight at 4 °C then incubated with secondary antibody (biotinylated rabbit anti-rat IgG, mouse absorbed, affinity purified, 1:200, Cat. no #BA-4001, Vector,) diluted in PBS supplemented with 5% normal rabbit serum for 45 min at RT, followed by incubation with streptavidin/HRP (1:500 dilution, cat #: P0397 Dako) diluted in PBS for 30 minutes in wet chamber at RT. Sections were counterstained with Gill’s hematoxylin solution (Sigma, cat #GHS316) and mounted with Glycerol-gelatin (cat #GG, Sigma). Data were collected using an Olympus VS120 Slide Scanner at 20x magnification. Quantification was performed using ImageJ.

### scRNA-seq library preparation

Whole bone marrow was isolated from long bones, hips, and vertebrae of four mice from each experimental group by crushing and then passaged through 70 μm cell strainers (Corning Cat. No. 352350 and 352340). Red cell lysis with ACK Lysing Buffer (Gibco Cat. No. 10492-01) was performed. Cells were washed, pooled and centrifuged and stained with PerCP-Cy™5.5 Mouse Lineage Antibody Cocktail (Cat. No. 561317, BD Pharmingen), anti-mouse CD117 (c-Kit) (Cat. No. 105818, Biolegend), and anti-mouse Ly-6A/E (Sca-1) (Cat. No. 108114, Biolegend) and sorted by FACS for Lineage negative, Sca1+, and c-Kit+ cells (LSK). Aortic arches from four mice from each experimental group were chopped into small pieces and digested using an enzyme mix containing 450 U/ml collagenase I (Sigma, cat# C0130), 125 U/ml collagenase XI (Sigma, cat# C7657), 60 U/ml DNase I (Sigma, cat# D4527) and 20 μM HEPES (ThermoFisher Scientific, cat# 15630106) for 45 min at 37°C and 750 rpm. Cells were washed, pooled, and centrifuged. LSK-sorted BM samples and aortic arch samples were stained with unique hashtag antibody–oligonucleotide conjugates against CD45.1 and CD45.2 (30 min) as per the manufacturer’s recommendations (TotalSeq, BioLegend). Samples were diluted to the manufacturer’s recommended concentration and immediately run on a 10X Chromium Controller after preparation with a 10X Chromium 3′ Library Preparation Kit (10X Genomics) to create the scRNA-seq libraries.

### 10X scRNA-seq and data preparation

Next-generation 150-nt paired-end sequencing was conducted on an Illumina NovaSeq 6000. The template switch oligonucleotide sequence was trimmed from the 5′ end and the poly(A) tail was trimmed from the 3′ end. Cell Ranger Count (10X Genomics) was used to filter low-quality reads and align to the mm9 mouse reference genome. The resulting files from the Cell Ranger pipeline were then converted to Seurat objects. Cells containing 10% or more reads mapping to the mitochondrial genome were filtered. Similarly, we filtered bone marrow cells with less than 500 genes and arch cells with less than 50 genes, and we filtered cells with more than 20,000 unique molecular identifiers. The doublet formation rate was set to 5%, based on standardized doublet rate expectations published by 10X Genomics^76^. Cells were determined to be i) CD45.1+ if the CD45.1 oligo count was >40 and the CD45.2 oligo count was <40, ii) CD45.2+ if the CD45.2 oligo count was >40 and the CD45.1 oligo count was <40, or negative for the expression of either CD45.1 or CD45.2 if the oligo counts were below 40 for both markers (non-hematopoietic cells). If both CD45.1 and CD45.2 oligo counts were > 40, then the cell was excluded as a doublet. Individual datasets were integrated with batch correction using a weighted nearest-neighbor approach. Supervised cell annotation was performed using the R package Azimuth using standard parameters and reference uniform manifold approximation and projection^77^ yielding expected cell types across all samples.

### scRNA-seq differential expression analysis and data visualization

To determine cell state differences in mice, differential expression was calculated using the FindMarkers function from the R package Seurat (v.4.1.1). Differences between cases and controls were quantified using the Wilcoxon rank-sum test. Genes with an absolute log_2_ fold change greater than 0.25 and adjusted *P*< 0.05 were considered statistically significant. Ribosomal and mitochondrial genes were excluded. Data visualization was performed using several R packages, including Seurat, and the Loupe Browser from 10x genomics.

### Western blotting

Macrophages were lysed in RIPA buffer containing protease inhibitor on ice for 10 min and then centrifuged at 14,000*g* for 5 min to generate protein lysates. Nuclear protein lysates were extracted using the kit from Abcam (Cat. No. ab113474). Protein concentration was determined by bicinchoninic assays and then mixed with 4× Laemmli buffer and heated at 95 °C for 5 min. Protein was separated by 4–12% gradient SDS–polyacrylamide gel electrophoresis and transferred onto nitrocellulose membranes. Then the membranes were blocked with 5% nonfat milk in Tris-buffered saline with 0.1% Tween 20 and immunostained with primary antibodies: anti-ATF3 (1:1,000 dilution, Cat. No. Ab207434, Abcam); and β-actin (1:1,000 dilution, Cat. No. 4970s, Cell Signaling Technology) at 4 °C overnight and detected using horseradish peroxidase (HRP)-conjugated secondary antibodies including HRP-linked anti-rabbit IgG (1:1,000 dilution, Cat. No. 7074S, Cell Signaling Technology).

### RNA-seq

RNA isolation was performed using the RNeasy Mini Kit (Qiagen), and RNA quality was assessed on a Fragment Analyzer (Advanced Analytical Technologies) − SS Total RNA 15nt. RNA-seq libraries were prepared using the Kapa mRNA HyperPrep Kit for Illumina (Roche) with Poly(A) selection according to the manufacturer’s instructions. Library quantification was examined on a Fragment Analyzer − HS NGS Fragment 1-6000bp and Qubit HS dsDNA Kit (Invitrogen). Libraries were pooled and sequenced to 150bp paired-end on the Illumina NovaSeq platform.

### RNA-Seq analysis

For expression analysis with bulk RNA-seq, reads were aligned to the mm9 revision of the mouse reference genome to which the sequences of the ERCC spike-in probes were added using hisat v2.1.0 ^78^ using default parameters. Aligned reads were converted into BAM format, sorted by name, and used as input for htseq-count ^79^ using parameters -i gene_id – stranded=reverse -m intersection-strict. The gene list quantified was built from RefSeq genes downloaded 2/1/2017 to which coordinates of the ERCC probes were added as distinct mock chromosomes. Counts were converted to transcripts per million (TPM) using the established protocol^80^. Specifically, all exons of all isoforms of each gene were collapsed using bedtools merge, and the sizes of these collapsed regions was used to calculate a per-sample normalization term, the for all genes of read length * number of reads / exon size; this normalization term was used accordingly, number of reads * read length / exon size * 1e6 / normalization term.

### ChIP-Seq

ChIP-seq was performed as previously described^81^.For each ChIP, 5 μg of H3K27ac antibody (Abcam) coupled to 2 μg of magnetic Dynabeads (Life Technologies) was added to 3 ml of sonicated nuclear extract from formaldehyde-fixed cells. Chromatin was immunoprecipitated overnight, cross-links were reversed, and DNA was purified by precipitation with phenol:chloroform:isoamyl alcohol. DNA pellets were resuspended in 25 μl of TE buffer. Illumina sequencing, library construction, and ChIP-seq analysis methods were previously described.

### CUT&RUN-Seq

CUT&RUN sequencing was performed as previously described with slight adaptations^82^. Concanavalin A (ConA) Conjugated Paramagnetic Beads (Fisher Scientific, #NC1526856) were initially activated using Bead Activation Buffer, containing 20 mM HEPES (pH 7.9), 10 mM KCl, 1 mM CaCl2, and 1 mM MnCl2. After activation, beads were washed and resuspended, preparing them for subsequent use. Cells were harvested and washed with RT Wash Buffer, and these cells were gently combined with the previously activated ConA beads to allow bead uptake by the cells. Next, antibodies for ATF3 or IgG, were added to the cell-bead complexes for specific targeting and incubated overnight at 4 degrees Celsius. The following day, the complexes were subjected to several washing and resuspension steps using Digitonin Buffer (Wash Buffer containing 0.01% Digitonin). In the final phase, targeted chromatin digestion and release were achieved using pAG-MNase (EpiCypher, #15-1016), alongside the addition of e. coli Spike-in DNA (EpiCypher, #18-1401). DNA purification was performed with the MinElute PCR purification kit (Qiagen, #28004). Illumina sequencing, library construction, and CUT&RUN-Seq analysis methods were previously described.

### ChIP-Seq and CUT&RUN-Seq analysis

For baseline analysis, raw ChIP-seq and CUT&RUN reads were aligned to the mm9 revision of the mouse reference genome using bowtie v 1.2.2 ^83^ in paired-end mode with parameters -k 1 - m 1 –best. Mated read-pairs were used to reconstruct sequence fragments using samtools v1.9 sort -n, bedtools v2.25.0 bamtobed -bedpe -i, and bedtools bedtobam -I ^84,85^. Coverage of 50bp bins with fragments was calculated by constructing a genome-wide set of bins with bedtools makewindows -w 50 and feeding these into bedtools intersect -c and normalizing for millions of fragments. Output bedGraph files were converted to bigwig for display using bedGraphToBigWig from UCSC tools^86^.

### Super-enhancers and assignment

Super-enhancers were identified using ROSE as previously described^87,88^. Briefly, H3K27ac and input control reads overlapping the ENCODE problematic regions list were identified and removed using bedtools intersect^89^. Remaining reads were used as input for two separate executions of MACS 1.4.1^90^ with parameters -p 1e-9, --keep-dup=auto and -p 1e-9 –keep- dup=all, and both using corresponding input control. The output peaks of the two MACS executions per sample were combined using bedtools merge and used as input for ROSE. ROSE parameters were -s 12500 -t 2000 -g mm9. Stitched enhancers were assigned to the single expressed transcript whose start site was nearest the center of the stitched enhancer, where expression was determined as being in the top 2/3 of genes ranked by promoter H3K27ac coverage, calculated by bamToGFF with parameters -m 1 -r -d.

### Heatmaps

For heatmap coverage display, raw paired-end reads were separately aligned in single-end mode to version mm9 of the mouse reference genome using bowtie v1.2.2 with parameters -k 1 -m 1 –best and -l set to the read length and combined into a single BAM file using samtools. Presumed PCR duplicates were removed using samtools rmdup. Peaks of ATF3 were called using MACS with parameters -p 1e-9 –keep-dup=auto and corresponding input control. The peaks identified independently from each of the three conditions were merged to a unified set using bedtools merge, and 4kb windows built off the midpoints of these collapsed peaks were used. Reads-per-million-normalized coverage matrices across the collapsed union of ATF3 peaks were generated using bamToGFF ^91^ with parameters -m 100 -r and ordered identically across the conditions using computed rowSums from the noted factor.

### Methylation quantification

Following the procedure adopted previously^23^, whole-genome bisulfite sequencing data were first processed with Trim Galore^92^ v0.6.10 with parameters -clip_r1 6 -clip_r2 6 -paired. Retained reads were analyzed using Bismark (0.24.2) ^93^ with parameters --non_directional --gzip -N 1 -- parallel 4 -p 4 --unmapped --nucleotide_coverage --score_min L,−10,−0.2 relative to the mm9 mouse reference genome. Bismark was also used to deduplicate events, and the methylation extractor was used to convert output to bedGraph format, which was converted to bigwig using UCSC tools. For per-CpG coverage tracks, the intersection of CpGs with >=2 reads in all five analyzed samples (wild-type DMSO, *Tet2*-/+ DMSO, *Tet2*-/- DMSO, *Tet2*-/- eltanexor, *Tet2*-/- eltanexor) were retained. To quantify methylation percentage in regions and compare methylation differences genome-wide, SeqMonk (v1.48.1)^94^ was used. Coverage files produced by Bismark were input, and read position probes were generated using requiring a minimum of 3 reads and spanning 50 valid positions across the five analyzed samples. Collective methylation of these probes was performed using the built-in “Bisulphite methylation over features” pipeline. Significant methylation differences of probes was defined as a >10% difference in percent methylation between conditions.

## QUANTIFICATION AND STATISTICAL ANALYSIS

### Data and Code Availability

Raw and processed data files were deposited to the NCBI GEO server under super-series GSE296964. Code written in R/python/Bash to perform analyses of ChIP-seq and CUT&RUN is available upon request.

### Additional Statistics

Data from the ChIP-seq experiments were analyzed as described above. Cell viability data were analyzed with two-way ANOVA followed by post-hoc t-test. Statistical significance was defined as a p-value <0.05. Data were analyzed with GraphPad Prism 9.4.0, and all error bars represent SD unless otherwise noted.

## References

1 Ahmad, H. & Jaiswal, S. Clonal haematopoiesis and atherosclerotic cardiovascular disease. Nat Rev Cardiol 20, 437–438 (2023). 10.1038/s41569-023-00882-2

2 Jaiswal, S. et al. Clonal Hematopoiesis and Risk of Atherosclerotic Cardiovascular Disease. N Engl J Med 377, 111–121 (2017). 10.1056/NEJMoa1701719

3 Weeks, L. D. et al. Prediction of risk for myeloid malignancy in clonal hematopoiesis. NEJM Evid 2 (2023). 10.1056/evidoa2200310

4 Tulstrup, M. et al. TET2 mutations are associated with hypermethylation at key regulatory enhancers in normal and malignant hematopoiesis. Nat Commun 12, 6061 (2021). 10.1038/s41467-021-26093-2

5 Moran-Crusio, K. et al. Tet2 loss leads to increased hematopoietic stem cell self-renewal and myeloid transformation. Cancer Cell 20, 11–24 (2011). 10.1016/j.ccr.2011.06.001

6 An, J. et al. Acute loss of TET function results in aggressive myeloid cancer in mice. Nat Commun 6, 10071 (2015). 10.1038/ncomms10071

7 Fuster, J. J. et al. Clonal hematopoiesis associated with TET2 deficiency accelerates atherosclerosis development in mice. Science 355, 842–847 (2017). 10.1126/science.aag1381

8 Colom Diaz, P. A., Mistry, J. J. & Trowbridge, J. J. Hematopoietic stem cell aging and leukemia transformation. Blood 142, 533–542 (2023). 10.1182/blood.2022017933

9 Wang, Y. et al. Tet2-mediated clonal hematopoiesis in nonconditioned mice accelerates age-associated cardiac dysfunction. JCI Insight 5 (2020). 10.1172/jci.insight.135204

10 Vromman, A. et al. Stage-dependent differential effects of interleukin-1 isoforms on experimental atherosclerosis. Eur Heart J 40, 2482–2491 (2019). 10.1093/eurheartj/ehz008

11 Liu, W. et al. Blockade of IL-6 signaling alleviates atherosclerosis in Tet2-deficient clonal hematopoiesis. Nat Cardiovasc Res 2, 572–586 (2023). 10.1038/s44161-023-00281-3

12 Lin, A. E. et al. Clonal Hematopoiesis of Indeterminate Potential With Loss of Tet2 Enhances Risk for Atrial Fibrillation Through Nlrp3 Inflammasome Activation. Circulation 149, 1419–1434 (2024). 10.1161/CIRCULATIONAHA.123.065597

13 Yeaton, A. et al. The Impact of Inflammation-Induced Tumor Plasticity during Myeloid Transformation. Cancer Discov 12, 2392–2413 (2022). 10.1158/2159-8290.CD-21-1146

14 Heyde, A. et al. Increased stem cell proliferation in atherosclerosis accelerates clonal hematopoiesis. Cell 184, 1348–1361 e1322 (2021). 10.1016/j.cell.2021.01.049

15 Jaiswal, S. & Libby, P. Author Correction: Clonal haematopoiesis: connecting ageing and inflammation in cardiovascular disease. Nat Rev Cardiol 17, 828 (2020). 10.1038/s41569-020-0414-8

16 Pardali, E., Dimmeler, S., Zeiher, A. M. & Rieger, M. A. Clonal hematopoiesis, aging, and cardiovascular diseases. Exp Hematol 83, 95–104 (2020). 10.1016/j.exphem.2019.12.006

17 Svensson, E. C. et al. TET2-Driven Clonal Hematopoiesis and Response to Canakinumab: An Exploratory Analysis of the CANTOS Randomized Clinical Trial. JAMA Cardiol 7, 521–528 (2022). 10.1001/jamacardio.2022.0386

18 Sano, S. et al. Tet2-Mediated Clonal Hematopoiesis Accelerates Heart Failure Through a Mechanism Involving the IL-1beta/NLRP3 Inflammasome. J Am Coll Cardiol 71, 875–886 (2018). 10.1016/j.jacc.2017.12.037

19 Zuriaga, M. A. et al. Colchicine prevents accelerated atherosclerosis in TET2-mutant clonal haematopoiesis. Eur Heart J (2024). 10.1093/eurheartj/ehae546

20 Jing, C. B. et al. Synthetic lethal targeting of TET2-mutant haematopoietic stem and progenitor cells by XPO1 inhibitors. Br J Haematol (2023). 10.1111/bjh.18667

21 Jing, C. B. et al. Synthetic lethal targeting of TET2-mutant haematopoietic stem and progenitor cells by XPO1 inhibitors. Br J Haematol 201, 489–501 (2023). 10.1111/bjh.18667

22 Jaiswal, S., Natarajan, P. & Ebert, B. L. Clonal Hematopoiesis and Atherosclerosis. N Engl J Med 377, 1401–1402 (2017). 10.1056/NEJMc1710381

23 McClatchy, J. et al. Clonal hematopoiesis related TET2 loss-of-function impedes IL1beta- mediated epigenetic reprogramming in hematopoietic stem and progenitor cells. Nat Commun 14, 8102 (2023). 10.1038/s41467-023-43697-y

24 Hopner, S. S. et al. LIGHT/LTbetaR signaling regulates self-renewal and differentiation of hematopoietic and leukemia stem cells. Nat Commun 12, 1065 (2021). 10.1038/s41467-021-21317-x

25 Culjkovic-Kraljacic, B. et al. Combinatorial targeting of nuclear export and translation of RNA inhibits aggressive B-cell lymphomas. Blood 127, 858–868 (2016). 10.1182/blood-2015-05-645069

26 Sommerville, L. J., Kelemen, S. E., Ellison, S. P., England, R. N. & Autieri, M. V. Increased atherosclerosis and vascular smooth muscle cell activation in AIF-1 transgenic mice fed a high-fat diet. Atherosclerosis 220, 45–52 (2012). 10.1016/j.atherosclerosis.2011.07.095

27 Seo, D. et al. Gene expression phenotypes of atherosclerosis. Arterioscler Thromb Vasc Biol 24, 1922–1927 (2004). 10.1161/01.ATV.0000141358.65242.1f

28 De Leon-Oliva, D. et al. AIF1: Function and Connection with Inflammatory Diseases. Biology (Basel) 12 (2023). 10.3390/biology12050694

29 Jadhav, K. & Zhang, Y. Activating transcription factor 3 in immune response and metabolic regulation. Liver Res 1, 96–102 (2017). 10.1016/j.livres.2017.08.001

30 Wu, J. et al. ATF3 and its emerging role in atherosclerosis: a narrative review. Cardiovasc Diagn Ther 12, 926–942 (2022). 10.21037/cdt-22-206

31 Hagen, M., Pangrazzi, L., Rocamora-Reverte, L. & Weinberger, B. Legend or Truth: Mature CD4(+)CD8(+) Double-Positive T Cells in the Periphery in Health and Disease. Biomedicines 11 (2023). 10.3390/biomedicines11102702

32 Takahashi, K. & Yamanaka, S. Induction of pluripotent stem cells from mouse embryonic and adult fibroblast cultures by defined factors. Cell 126, 663–676 (2006). 10.1016/j.cell.2006.07.024

33 Feinberg, M. W. et al. Kruppel-like factor 4 is a mediator of proinflammatory signaling in macrophages. J Biol Chem 280, 38247–38258 (2005). 10.1074/jbc.M509378200

34 Kim, A. D. et al. Correction to: Myeloid-specific deletion of chitinase-3-like 1 protein ameliorates murine diet-induced steatohepatitis progression. J Mol Med (Berl) 101, 1627 (2023). 10.1007/s00109-023-02388-3

35 Guo, Q. et al. NF-kappaB in biology and targeted therapy: new insights and translational implications. Signal Transduct Target Ther 9, 53 (2024). 10.1038/s41392-024-01757-9

36 Lawrence, T. The nuclear factor NF-kappaB pathway in inflammation. Cold Spring Harb Perspect Biol 1, a001651 (2009). 10.1101/cshperspect.a001651

37 Liu, T., Zhang, L., Joo, D. & Sun, S. C. NF-kappaB signaling in inflammation. Signal Transduct Target Ther 2, 17023- (2017). 10.1038/sigtrans.2017.23

38 Martin-Ventura, J. L. et al. Increased CD74 expression in human atherosclerotic plaques: contribution to inflammatory responses in vascular cells. Cardiovasc Res 83, 586–594 (2009). 10.1093/cvr/cvp141

39 de Aguiar, M. F. et al. Monocyte subsets and monocyte-related chemokines in Takayasu arteritis. Sci Rep 13, 2092 (2023). 10.1038/s41598-023-29369-3

40 Chang, T. T. et al. Inhibition of CCL7 improves endothelial dysfunction and vasculopathy in mouse models of diabetes mellitus. Sci Transl Med 16, eadn1507 (2024). 10.1126/scitranslmed.adn1507

41 Bennett, M. R., Sinha, S. & Owens, G. K. Vascular Smooth Muscle Cells in Atherosclerosis. Circ Res 118, 692–702 (2016). 10.1161/CIRCRESAHA.115.306361

42 Mussbacher, M. et al. More than Just a Monolayer: the Multifaceted Role of Endothelial Cells in the Pathophysiology of Atherosclerosis. Curr Atheroscler Rep 24, 483–492 (2022). 10.1007/s11883-022-01023-9

43 Kong, P. et al. Inflammation and atherosclerosis: signaling pathways and therapeutic intervention. Signal Transduct Target Ther 7, 131 (2022). 10.1038/s41392-022-00955-7

44 Muhl, L. et al. A single-cell transcriptomic inventory of murine smooth muscle cells. Dev Cell 57, 2426–2443 e2426 (2022). 10.1016/j.devcel.2022.09.015

45 Dong, R., Jiang, G., Tian, Y. & Shi, X. Identification of immune-related biomarkers and construction of regulatory network in patients with atherosclerosis. BMC Med Genomics 15, 245 (2022). 10.1186/s12920-022-01397-4

46 Vaittinen, M. et al. MFAP5 is related to obesity-associated adipose tissue and extracellular matrix remodeling and inflammation. Obesity (Silver Spring) 23, 1371–1378 (2015). 10.1002/oby.21103

47 Kiss, M. G. et al. Cell-autonomous regulation of complement C3 by factor H limits macrophage efferocytosis and exacerbates atherosclerosis. Immunity 56, 1809–1824 e1810 (2023). 10.1016/j.immuni.2023.06.026

48 Bondareva, O. et al. Single-cell profiling of vascular endothelial cells reveals progressive organ-specific vulnerabilities during obesity. Nat Metab 4, 1591–1610 (2022). 10.1038/s42255-022-00674-x

49 Miao, G. et al. Vascular smooth muscle cell c-Fos is critical for foam cell formation and atherosclerosis. Metabolism 132, 155213 (2022). 10.1016/j.metabol.2022.155213

50 Hoeflich, A., David, R. & Hjortebjerg, R. Current IGFBP-Related Biomarker Research in Cardiovascular Disease-We Need More Structural and Functional Information in Clinical Studies. Front Endocrinol (Lausanne) 9, 388 (2018). 10.3389/fendo.2018.00388

51 Glickman, J. W. et al. Cross-sectional study of blood biomarkers of patients with moderate to severe alopecia areata reveals systemic immune and cardiovascular biomarker dysregulation. J Am Acad Dermatol 84, 370–380 (2021). 10.1016/j.jaad.2020.04.138

52 Sjaarda, J. et al. Blood CSF1 and CXCL12 as Causal Mediators of Coronary Artery Disease. J Am Coll Cardiol 72, 300–310 (2018). 10.1016/j.jacc.2018.04.067

53 Pickett, J. R., Wu, Y., Zacchi, L. F. & Ta, H. T. Targeting endothelial vascular cell adhesion molecule-1 in atherosclerosis: drug discovery and development of vascular cell adhesion molecule-1-directed novel therapeutics. Cardiovasc Res 119, 2278–2293 (2023). 10.1093/cvr/cvad130

54 Rasmussen, K. D. et al. Loss of TET2 in hematopoietic cells leads to DNA hypermethylation of active enhancers and induction of leukemogenesis. Genes Dev 29, 910–922 (2015). 10.1101/gad.260174.115

55 Hnisz, D. et al. Super-enhancers in the control of cell identity and disease. Cell 155, 934–947 (2013). 10.1016/j.cell.2013.09.053

56 Parker, S. C. et al. Chromatin stretch enhancer states drive cell-specific gene regulation and harbor human disease risk variants. Proc Natl Acad Sci U S A 110, 17921–17926 (2013). 10.1073/pnas.1317023110

57 Rodriguez-Martinez, J. A., Reinke, A. W., Bhimsaria, D., Keating, A. E. & Ansari, A. Z. Combinatorial bZIP dimers display complex DNA-binding specificity landscapes. Elife 6 (2017). 10.7554/eLife.19272

58 Zhou, J. et al. Anti-inflammatory Activity of MTL-CEBPA, a Small Activating RNA Drug, in LPS-Stimulated Monocytes and Humanized Mice. Mol Ther 27, 999–1016 (2019). 10.1016/j.ymthe.2019.02.018

59 Skene, P. J. & Henikoff, S. An efficient targeted nuclease strategy for high-resolution mapping of DNA binding sites. Elife 6 (2017). 10.7554/eLife.21856

60 Gold, E. S. et al. ATF3 protects against atherosclerosis by suppressing 25- hydroxycholesterol-induced lipid body formation. J Exp Med 209, 807–817 (2012). 10.1084/jem.20111202

61 Labzin, L. I. et al. ATF3 Is a Key Regulator of Macrophage IFN Responses. J Immunol 195, 4446–4455 (2015). 10.4049/jimmunol.1500204

62 Sha, H., Zhang, D., Zhang, Y., Wen, Y. & Wang, Y. ATF3 promotes migration and M1/M2 polarization of macrophages by activating tenascin-C via Wnt/beta-catenin pathway. Mol Med Rep 16, 3641–3647 (2017). 10.3892/mmr.2017.6992

63 Ito, S. et al. Role of Tet proteins in 5mC to 5hmC conversion, ES-cell self-renewal and inner cell mass specification. Nature 466, 1129–1133 (2010). 10.1038/nature09303

64 Kribelbauer, J. F. et al. Quantitative Analysis of the DNA Methylation Sensitivity of Transcription Factor Complexes. Cell Rep 19, 2383–2395 (2017). 10.1016/j.celrep.2017.05.069

65 Fidler, T. P. et al. Suppression of IL-1beta promotes beneficial accumulation of fibroblast-like cells in atherosclerotic plaques in clonal hematopoiesis. Nat Cardiovasc Res 3, 60–75 (2024). 10.1038/s44161-023-00405-9

66 Wang, S. et al. Combining selective inhibitors of nuclear export (SINEs) with chimeric antigen receptor (CAR) T cells for CD19-positive malignancies. Oncol Rep 46 (2021). 10.3892/or.2021.8121

67 Hing, Z. A. et al. Next-generation XPO1 inhibitor shows improved efficacy and in vivo tolerability in hematological malignancies. Leukemia 30, 2364–2372 (2016). 10.1038/leu.2016.136

68 Fernandez, D. M. et al. Single-cell immune landscape of human atherosclerotic plaques. Nat Med 25, 1576–1588 (2019). 10.1038/s41591-019-0590-4

69 Nguyen, C. T., Kim, E. H., Luong, T. T., Pyo, S. & Rhee, D. K. TLR4 mediates pneumolysin-induced ATF3 expression through the JNK/p38 pathway in Streptococcus pneumoniae- infected RAW 264.7 cells. Mol Cells 38, 58–64 (2015). 10.14348/molcells.2015.2231

70 Wang, Y. et al. Smooth Muscle Cells Contribute the Majority of Foam Cells in ApoE (Apolipoprotein E)-Deficient Mouse Atherosclerosis. Arterioscler Thromb Vasc Biol 39, 876–887 (2019). 10.1161/ATVBAHA.119.312434

71 Libby, P. Inflammation in atherosclerosis. Arterioscler Thromb Vasc Biol 32, 2045–2051 (2012). 10.1161/ATVBAHA.108.179705

72 Zhang, X., Sessa, W. C. & Fernandez-Hernando, C. Endothelial Transcytosis of Lipoproteins in Atherosclerosis. Front Cardiovasc Med 5, 130 (2018). 10.3389/fcvm.2018.00130

73 Pepin, M. E. & Gupta, R. M. The Role of Endothelial Cells in Atherosclerosis: Insights from Genetic Association Studies. Am J Pathol 194, 499–509 (2024). 10.1016/j.ajpath.2023.09.012

74 Singh, S. & Torzewski, M. Fibroblasts and Their Pathological Functions in the Fibrosis of Aortic Valve Sclerosis and Atherosclerosis. Biomolecules 9 (2019). 10.3390/biom9090472

75 Rasmussen, K. D. et al. TET2 binding to enhancers facilitates transcription factor recruitment in hematopoietic cells. Genome Res 29, 564–575 (2019). 10.1101/gr.239277.118

76. Genomics, x. Technical Note CG000148 – Resolving Cell Types as a Function of Read Depth and Cell Number, <https://assets.ctfassets.net/an68im79xiti/6gDArDPBTOg4IIkYEO2Sis/803be2286bba5ca67f353e6baf68d276/CG000148_10x_Technical_Note_Resolving_Cell_Types_as_Function_of_Read_Depth_Cell_Number_RevA.pdf> (2018).

77. Satija. Azimuth Reference, <https://zenodo.org/records/4546839#.Ytlg9uzMLS4> (2021).

78 Kim, D., Langmead, B. & Salzberg, S. L. HISAT: a fast spliced aligner with low memory requirements. Nat Methods 12, 357–360 (2015). 10.1038/nmeth.3317

79 Anders, S., Pyl, P. T. & Huber, W. HTSeq--a Python framework to work with high-throughput sequencing data. Bioinformatics 31, 166–169 (2015). 10.1093/bioinformatics/btu638

80 Wagner, G. P., Kin, K. & Lynch, V. J. Measurement of mRNA abundance using RNA-seq data: RPKM measure is inconsistent among samples. Theory Biosci 131, 281–285 (2012). 10.1007/s12064-012-0162-3

81 Wong, R. W. J. et al. Enhancer profiling identifies critical cancer genes and characterizes cell identity in adult T-cell leukemia. Blood 130, 2326–2338 (2017). 10.1182/blood-2017-06-792184

82 Skene, P. J., Henikoff, J. G. & Henikoff, S. Targeted in situ genome-wide profiling with high efficiency for low cell numbers. Nat Protoc 13, 1006–1019 (2018). 10.1038/nprot.2018.015

83 Langmead, B., Trapnell, C., Pop, M. & Salzberg, S. L. Ultrafast and memory-efficient alignment of short DNA sequences to the human genome. Genome Biol 10, R25 (2009). 10.1186/gb-2009-10-3-r25

84 Li, H. et al. The Sequence Alignment/Map format and SAMtools. Bioinformatics 25, 2078–2079 (2009). 10.1093/bioinformatics/btp352

85 Quinlan, A. R. & Hall, I. M. BEDTools: a flexible suite of utilities for comparing genomic features. Bioinformatics 26, 841–842 (2010). 10.1093/bioinformatics/btq033

86 Fujita, P. A. et al. The UCSC Genome Browser database: update 2011. Nucleic Acids Res 39, D876–882 (2011). 10.1093/nar/gkq963

87 Whyte, W. A. et al. Master transcription factors and mediator establish super-enhancers at key cell identity genes. Cell 153, 307–319 (2013). 10.1016/j.cell.2013.03.035

88 Loven, J. et al. Selective inhibition of tumor oncogenes by disruption of super-enhancers. Cell 153, 320–334 (2013). 10.1016/j.cell.2013.03.036

89 Amemiya, H. M., Kundaje, A. & Boyle, A. P. The ENCODE Blacklist: Identification of Problematic Regions of the Genome. Sci Rep 9, 9354 (2019). 10.1038/s41598-019-45839-z

90 Zhang, Y. et al. Model-based analysis of ChIP-Seq (MACS). Genome Biol 9, R137 (2008). 10.1186/gb-2008-9-9-r137

91. Bradner, J. E. Bradner Lab Pipeline, <https://github.com/BradnerLab/pipeline> (2013).

92. Krueger, F. *Trim Galore*, <https://github.com/FelixKrueger/TrimGalore> (

93 Krueger, F. & Andrews, S. R. Bismark: a flexible aligner and methylation caller for Bisulfite- Seq applications. Bioinformatics 27, 1571–1572 (2011). 10.1093/bioinformatics/btr167

94. S, A. SeqMonk, <https://github.com/s-andrews/SeqMonk> (

