## Extended Data Figures for "Multitargeted Reduction of Inflammation and Atherosclerosis in *Tet2*-deficient CHIP via XPO1 Inhibition and Atf3 restoration"

### Extended Data Figure 1

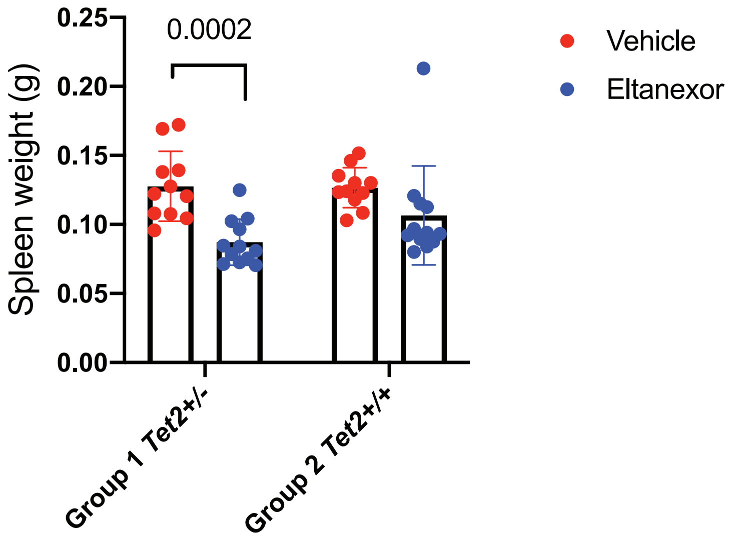

### Extended Data Figure 2

**a**

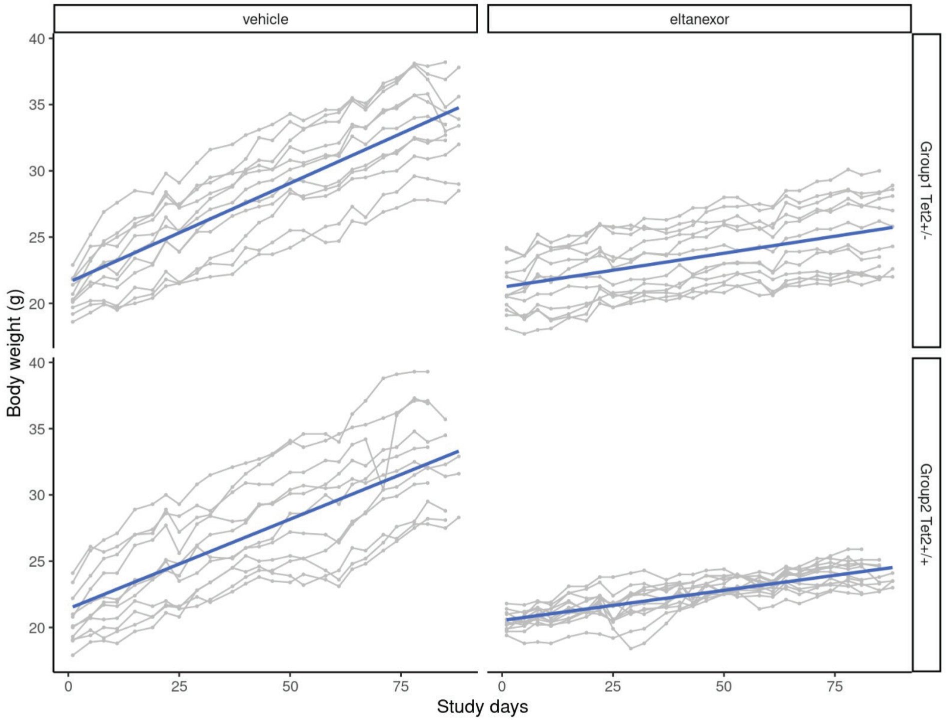


**b**

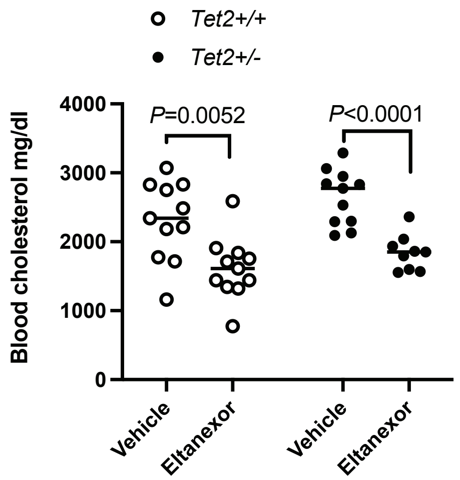

### Extended Data Figure 3

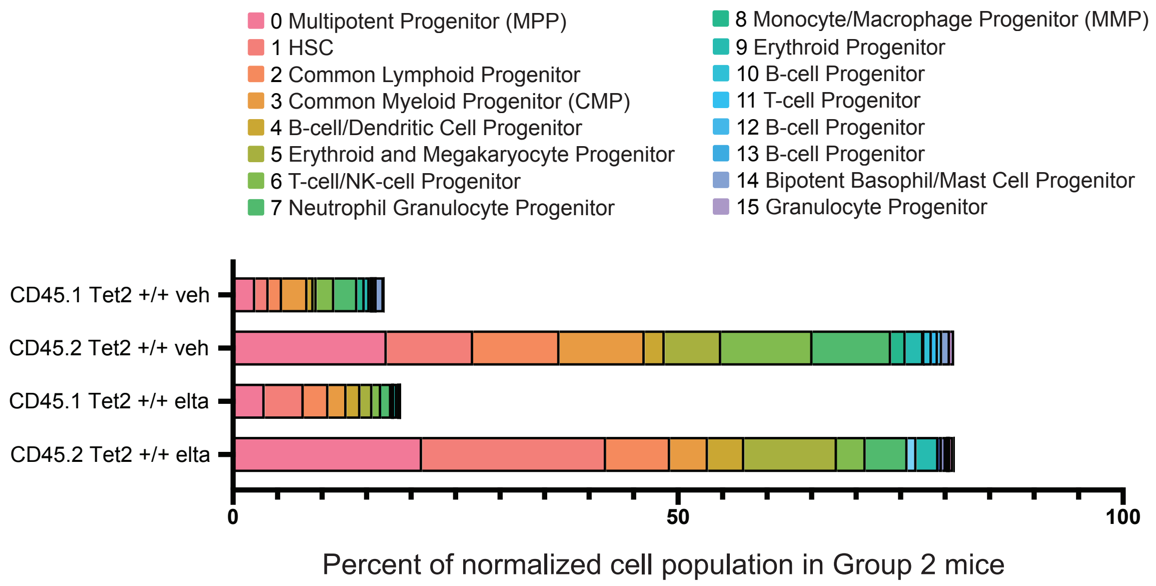

### Extended Data Figure 4

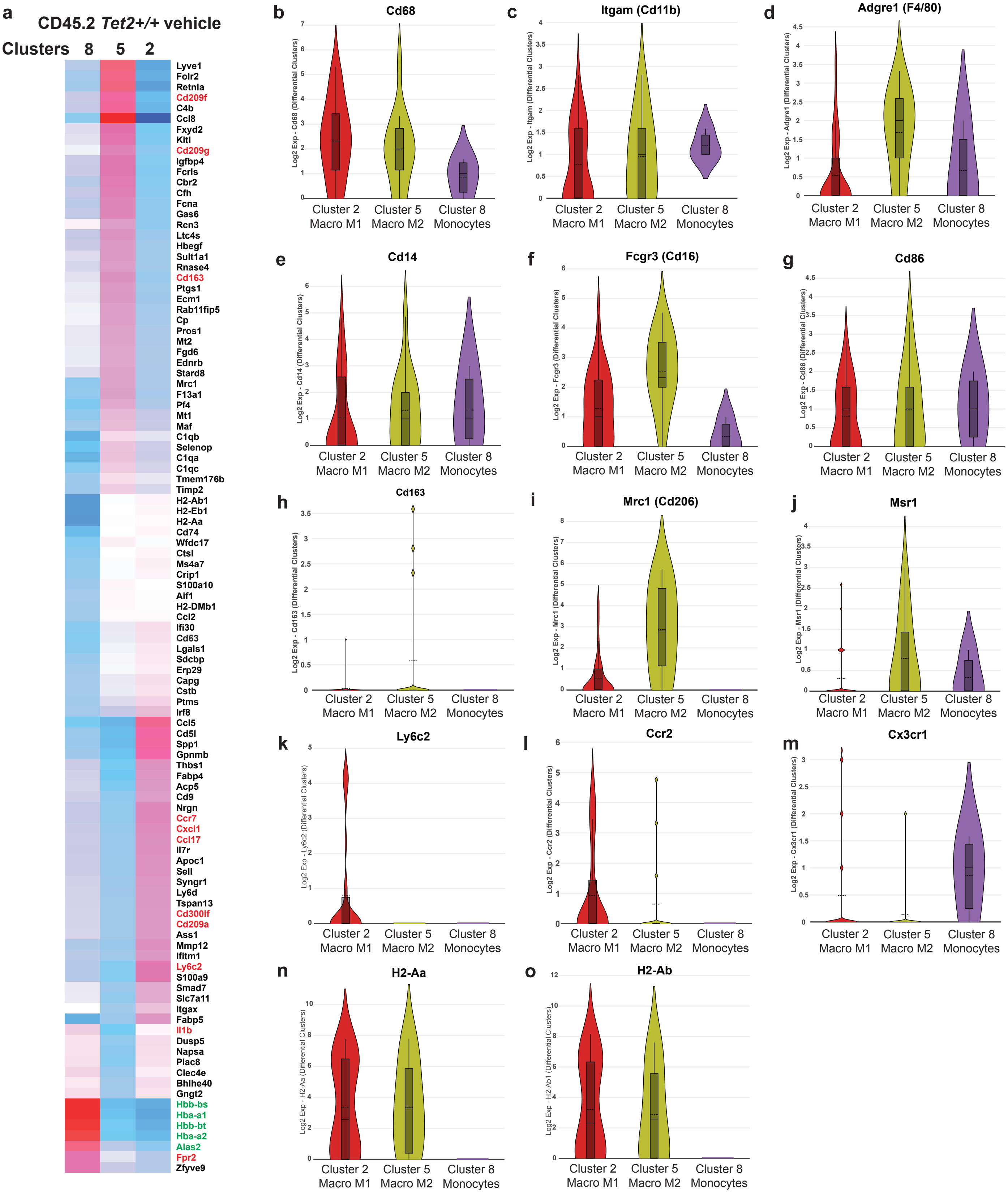

#### Extended Data Figure 5

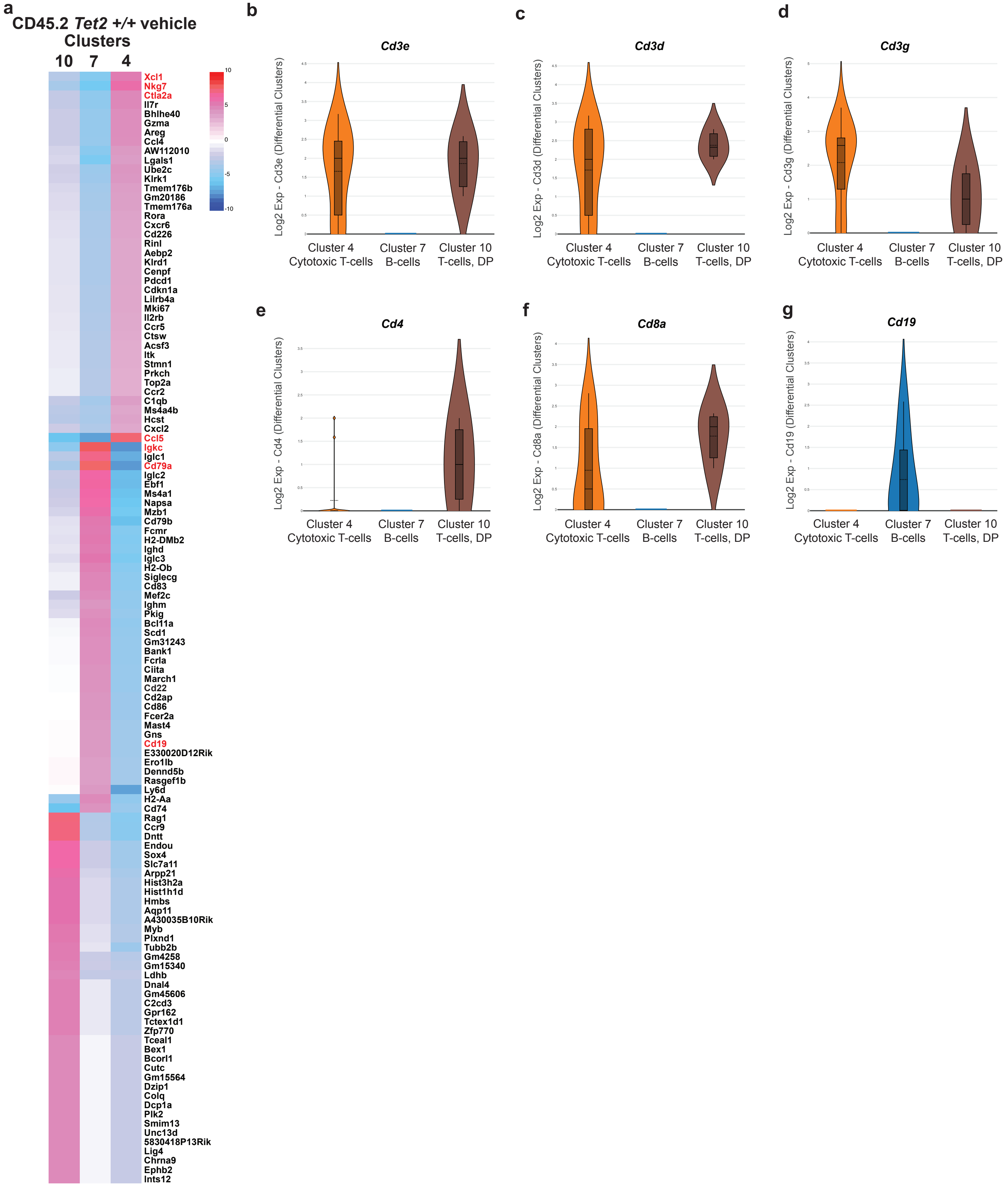

### Extended Data Figure 6

a

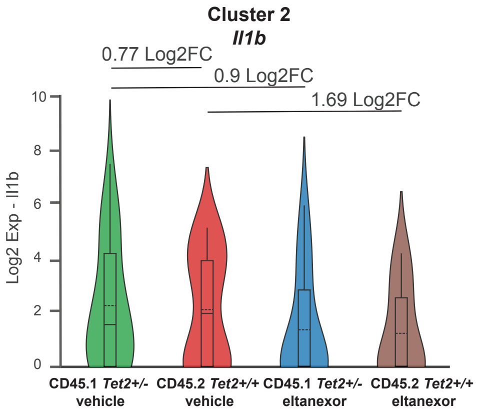


b

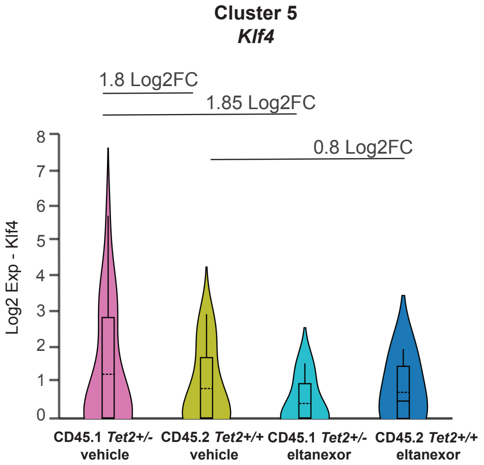

### Extended Data Figure 7

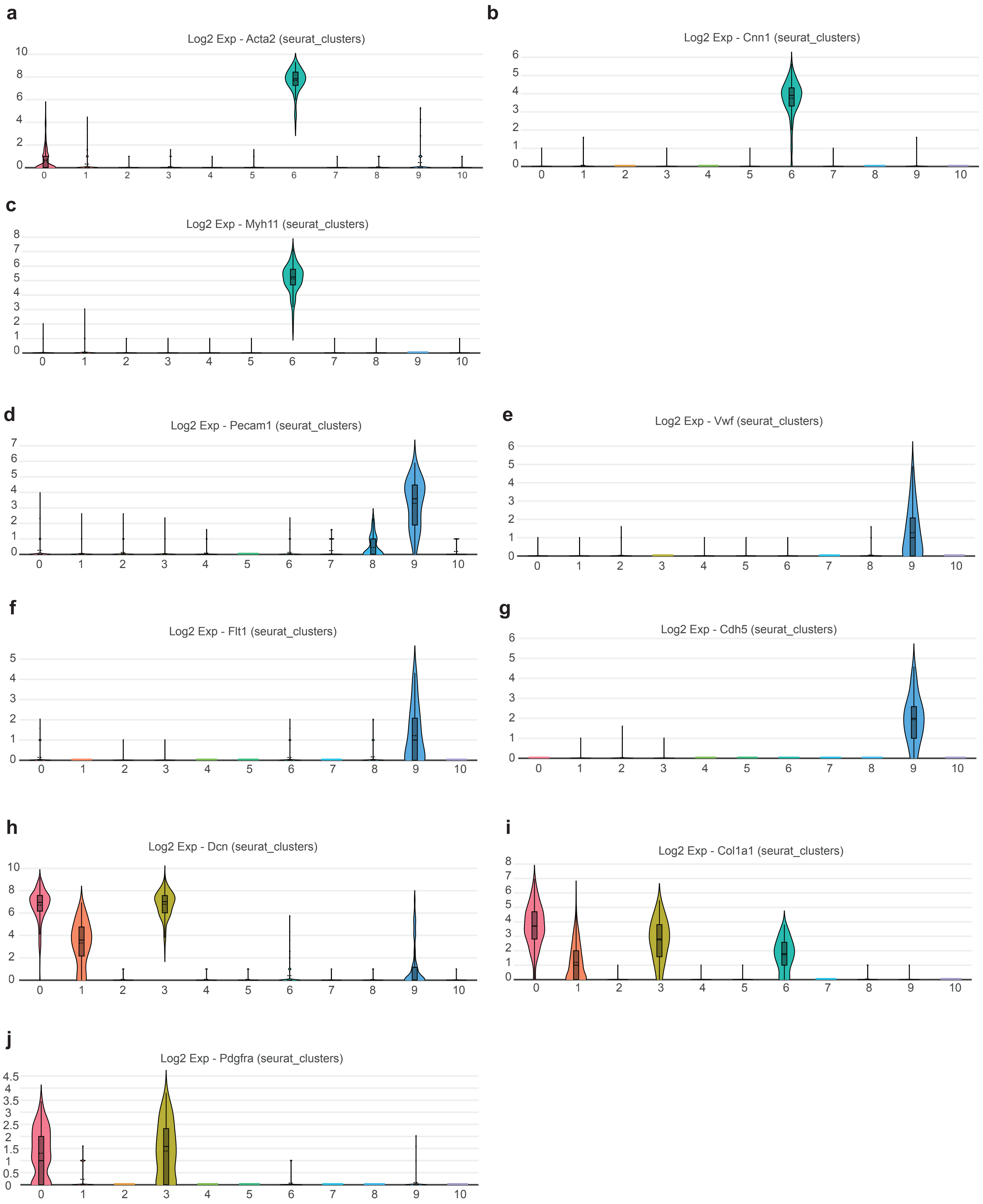

### Extended Data Figure 8

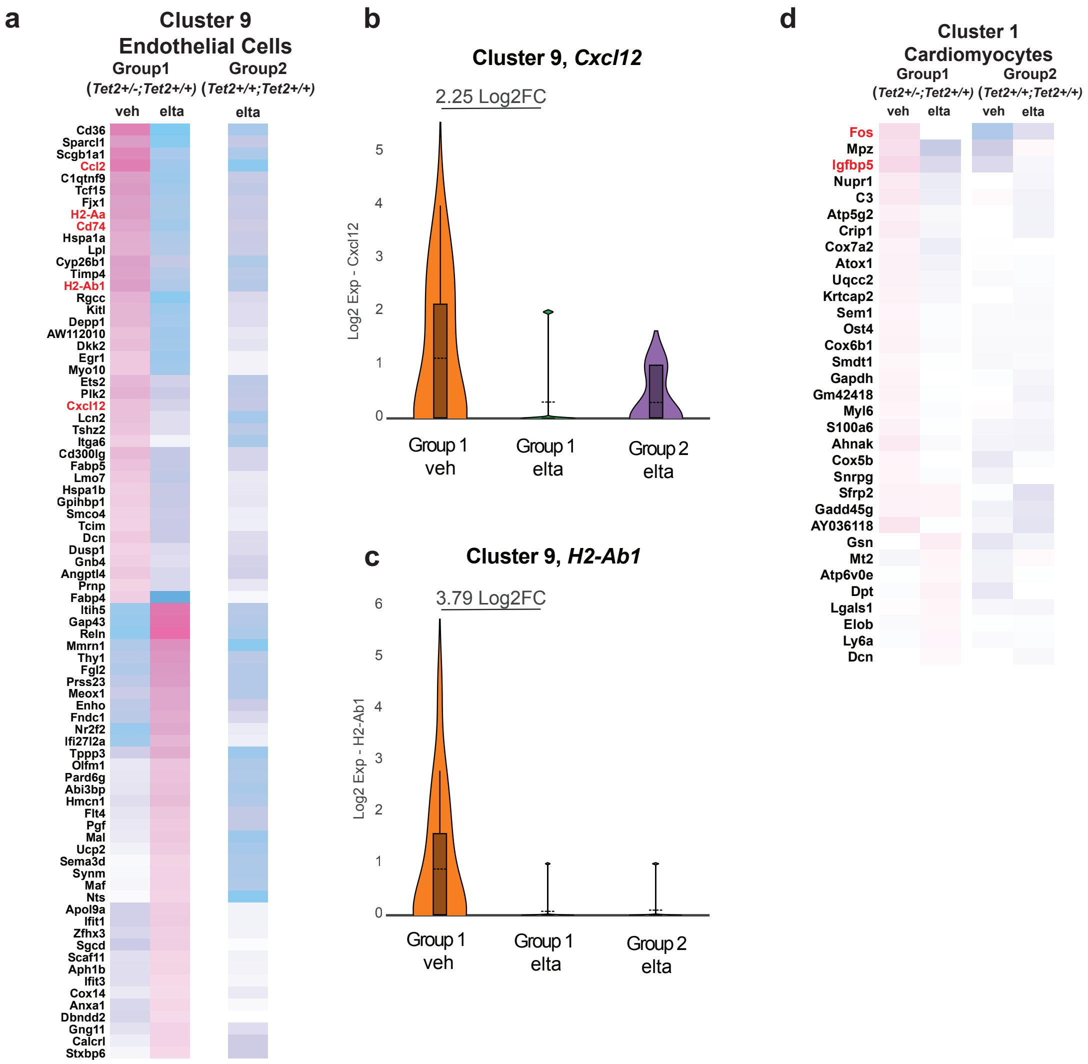

#### Extended Data Figure 9

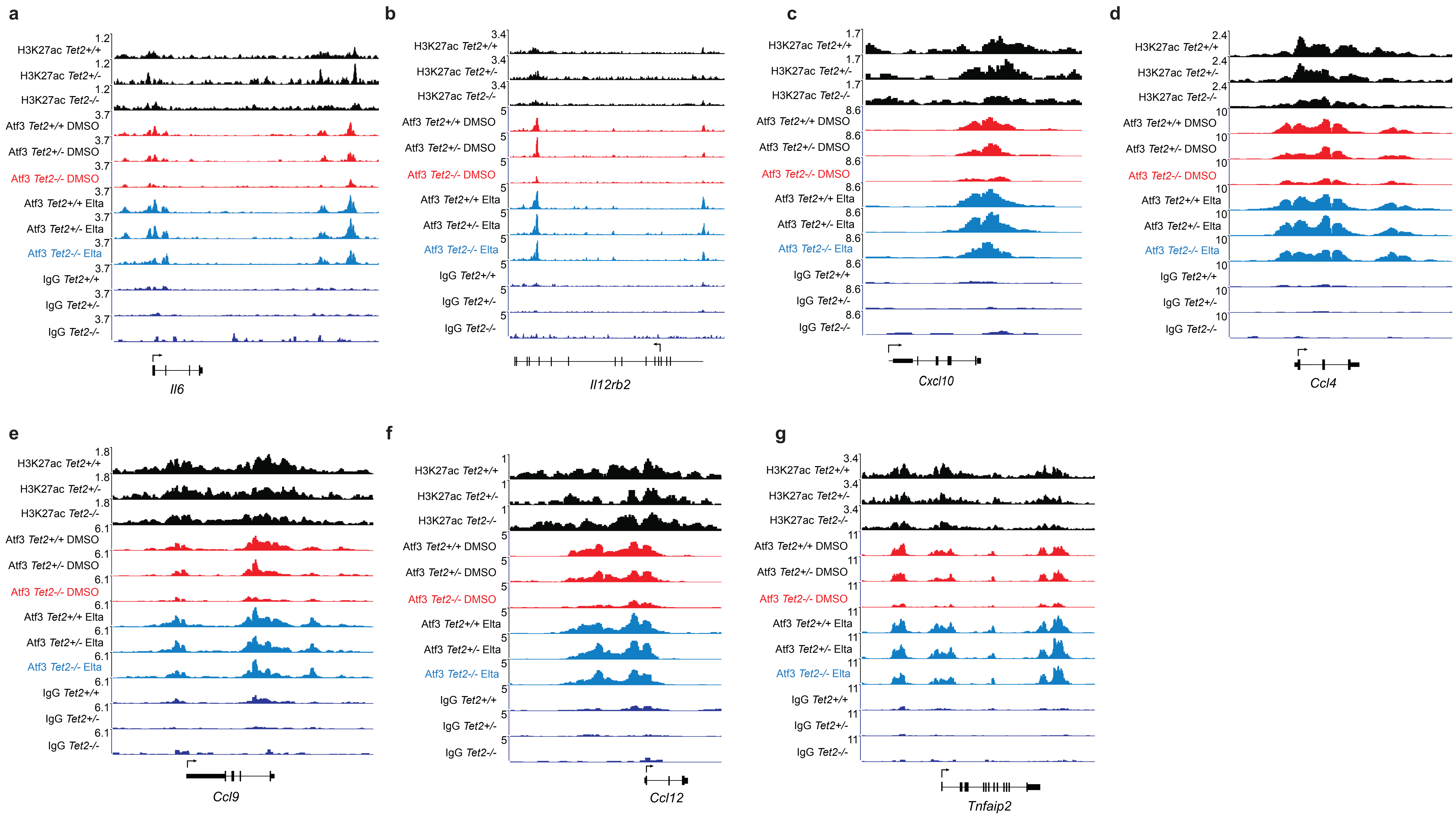

**Extended Data Figure 1. Eltanexor selectively reduces spleen weight in *Ldlr-/-* mice injected with *Tet2+/-* bone marrow cells.** Spleen weights of vehicle-treated (red) and eltanexor-treated (blue) mice injected with *Tet2+/-*;CD45.1+ bone marrow cells (Group 1) or *Tet2+/+*;CD45.1+ bone marrow cells (Group 2). P-values obtained by unpaired t-test.

**Extended Data Figure 2. Eltanexor-treated mice gain less weight on a high cholesterol diet. a,** Mean body weight of Group 2 *Ldlr-/-* mice injected with a mix of *Tet2+/+*;CD45.1+ and *Tet2+/+*;CD45.2+ cells consuming a high cholesterol diet over the course of 12 weeks. Vehicle treated mice are indicated in red and eltanexor treated mice are indicated in blue. **b,** Mean body weight of Group 1 *Ldlr-/-* mice injected with a mix of *Tet2+/-*;CD45.1+ and *Tet2+/+*;CD45.2+ cells consuming a high cholesterol diet over the course of 12 weeks. Vehicle treated mice are indicated in red and eltanexor treated mice are indicated in blue. **c**, Blood cholesterol levels in mg/dl of vehicle-treated and eltanexor-treated mice injected with *Tet2+/+*;CD45.1+ bone marrow (Group 2) or *Tet2+/-*;CD45.1+ bone marrow (Group 1) after 12 weeks of a high cholesterol diet. P-values obtained by unpaired t-test.

**Extended Data Figure 3. Single-cell CITEseq results for LSK-sorted bone marrow cells after 12 weeks of *Tet2+/+;*CD45.2+ bone marrow cells compared to *Tet2+/+*;CD45.1+ bone marrow cells of Group 2 vehicle- and eltanexor-treated mice.** Frequency bar chart comparing the proportions of cells within 16 UMAP clusters of LSK-sorted bone marrow cels at 12 weeks comparing *Tet2+/+*;CD45.1+ and *Tet2+/+*;CD45.2+ cell populations in mice treated with vehicle or eltanexor.

**Extended Data Figure 4. Identification of monocytes and macrophage subsets. a**, Heatmap of most differentially expressed genes between *Tet2+/+;*CD45.2+ cells of Clusters 8 (monocytes), 5 (M2 macrophages), and 2 (M1 macrophages) in vehicle treated mice. **b-o**, Violin plots of scRNA-seq data from *Tet2+/+*;CD45.2+ cells from vehicle-treated mice showing expression of macrophage and monocyte markers in Clusters 2, 5, and 8 for *Cd68* (b) *Cd11b* (c), *F4/80* (d), *Cd14* (e), *Cd16* (f), *Cd86* (g), *Cd163* (h), *Cd206* (i), *Msr1* (j), *Ly6c2* (k), *Ccr2* (l), *Cx3cr1* (m), *H2-Aa* (n), *H2-Ab* (o).

**Extended Data Figure 5. Identification of B- and T-cells. a**, Heatmap of most differentially expressed genes between *Tet2+/+*;CD45.2+ cells of Clusters 10 (double positive T-cells), 7 (B-cells), and 4 (cytotoxic T-cells) in vehicle treated mice. **b-g**, Violin plots of scRNA-seq data from

*Tet2+/+*;CD45.2+ cells from vehicle treated mice showing expression of T- and B-cell markers for *Cd3e* (b), *Cd3d* (c), *Cd3e* (d), *Cd4* (e), *Cd8a* (f), *Cd19* (g).

**Extended Data Figure 6. Eltanexor treatment decreases the expression levels of inflammatory mediators in M1 macrophages**. **a-c**, Violin plots of scRNA-seq data from *Tet2+/-;*CD45.1+ and *Tet2+/+;*CD45.2+ Cluster 2 M1 macrophages and Cluster 5 M2 macrophages in mice treated with vehicle or eltanexor for *Il1b* (a), *Klf4* (b).

**Extended Data Figure 7. Identification of non-hematopoietic cells.** Violin plots of scRNA-seq data from non-hematopoietic cells from vehicle treated mice showing expression of Cluster 6 smooth muscle cell markers *Acta2* (a), *Cnn1* (b) and *Myh11* (c), Cluster 9 endothelial cell markers *Pecam1* (d), *Vwf* (e), *Flt1* (f), *Cdh5* (g), and Cluster 3 fibroblast markers *Dcn* (h), *Col1a1* (i), and *Pdgfra* (j). Cluster 0 fibroblasts also express *Acta2* (a) and have been identified as myofibroblasts.

**Extended Data Figure 8. Eltanexor reduces the expression of inflammatory genes in endothelial cells. a,** Heatmap of differentially expressed genes in endothelial cells from eltanexor- and vehicle-treated control mice injected with*Tet2+/-;*CD45.1+ bone marrow cells (Group 1) relative to endothelial cells from eltanexor treated mice injected with *Tet2+/+;*CD45.1+ bone marrow cells (Group 2). Note: endothelial cell numbers recovered from vehicle treated Group 2 mice were not sufficient for analysis. **b-c**, Violin plots of scRNA-seq data from endothelial cells from vehicle- and eltanexor-treated Group 1 mice and eltanexor-treated Group 2 mice showing expression of *Cxcl12* (b) and *H2-Ab1* (c). **d**, Heatmap of differentially expressed genes in cardiomyocytes from eltanexor treated and vehicle control mice injected with *Tet2+/-;*CD45.1+ bone marrow cells (Group 1) relative to endothelial cells from mice injected with *Tet2+/+;*CD45.1+ bone marrow cells (Group 2).

**Extended Data Figure 9. Atf3 binding to the enhancers of multiple inflammatory genes is reduced in *Tet2*-mutant BMDMs. a-h,** CUT&RUN sequencing alignment tracks for Atf3 and IgG (control) in DSMO control- or eltanexor (Elta)-treated BMDM from *Tet2+/+, Tet2+/-* or *Tet2-/-* mice, overlaid with H3K27ac ChIP-seq at the *Il6* locus (a), the *Il12rb2* locus (b), the *Cxcl10* locus (c), the *Ccl4* locus (d), the *Ccl9* locus (e), the *Ccl12* locus (f), and the *Tnfaip2* locus (g). The Atf3 Cut-and-Run track for DMSO-treated, *Tet2-/-* BMDMs is highlighted in red font, and the

Atf3 Cut-and-Run track for eltanexor-treated, *Tet2-/-* BMDMs is highlighted in blue font in each panel.
